# Once yearly cell-based therapy for sustained and dose tunable delivery of monoclonal antibodies

**DOI:** 10.64898/2026.05.19.726224

**Authors:** Cody Fell, Anthony E. Davis, Shalini Pandey, Michael T. Guinn, Zeshi Wang, Jonathon DeBonis, Chancellor Smith, Nathan Brown, Danna Murungi, Yeonju Kim, Imaan Mohandessi, Patrick Bednarz, Amir Ardeshir, Erin M. Haupt, Samuel I. Cuevas, Christy L. Lavine, Michael S. Seaman, Oleg Igoshin, Ravi K. Ghanta, Michael R. Diehl, Omid Veiseh

## Abstract

Over 200 monoclonal antibodies (mAbs) are approved for clinical use, yet their therapeutic potential is constrained by dependence on repeated injections or infusions that drive non-adherence, limit access in low-resource settings, and generate peak–trough pharmacokinetics linked to adverse effects and reduced efficacy. Here, we developed an immunomodulatory encapsulated cell-based ‘biologics factory’ that overcomes mAb instability, immunogenicity, and the fibrotic foreign body response that have limited previous approaches, enabling continuous *in situ* production of therapeutic antibodies from a single administration. Screening chemically modified alginate biomaterials in immunocompetent mice identified a lead immunomodulatory alginate formulation that sustains stable serum titers of the HIV-neutralizing mAb 3BNC117 for one year. Single-cell RNA sequencing revealed that this formulation promotes a local anti-inflammatory, pro-resolving immune niche that attenuates fibrosis. The platform’s versatility was demonstrated by production of thirteen diverse mAbs from an allogeneic cell chassis, with sustained *in vivo* delivery of a subset including ipilimumab, pembrolizumab, adalimumab, and PGT121. Integration into a retrievable macrodevice enabled on-demand therapeutic termination and re-implantation for dose-proportional tuning. In a non-human primates, subcutaneous implantation maintained stable ipilimumab titers for over six months with no detectable toxicity, anti-drug antibodies, or adverse events, and dose-dependent exposure was confirmed across a three-dose escalation. These results demonstrate a clinically translatable platform offering a practical strategy to replace frequent injections with single-administration therapy.

## Main

There are currently more than 212 monoclonal antibodies approved for clinical use worldwide, which are reshaping outcomes across many disease indication areas, including autoimmunity, infectious disease, and oncology^1^. Yet, current delivery methods still depend on frequent high-dose intravenous (IV) infusions and subcutaneous or intramuscular (SC/IM) bolus injections. These regimens generate peak–trough pharmacokinetics linked to off-target effects at peaks, reduced efficacy at troughs, high cost and logistical burden, and reduced adherence, especially in patients with chronic diseases^2–4^. A broadly applicable solution for mAb delivery would provide (i) dose adjustability to meet patient-specific pharmacokinetic–pharmacodynamic (PK/PD) needs, (ii) long-term sustained delivery on the scale of months to years, and (iii) the option for rapid termination of therapeutic delivery to mitigate adverse events or accommodate changes in therapy.

Several strategies aim to extend mAb exposure beyond conventional bolus dosing. Polymeric slow release formulations and mRNA–lipid nanoparticle approaches can reduce dose frequency or enable transient *in vivo* expression, respectively, but face challenges including mAb instability, depot-induced aggregation, liver-tropic biodistribution, immunogenicity upon redosing, and difficulty in rapidly terminating exposure^5–9^. These approaches improve convenience and access beyond IV infusions, but they do not yet meet the combined requirements of durable, tunable, and reversible systemic mAb delivery for chronic indications. Gene therapy strategies like AAV-mediated transduction and B-/plasma-cell engineering have enabled durable year-long expression but introduce distinct control and safety trade-offs. AAV therapies are subject to anti-capsid immunity which limits redosing, and clinically meaningful down-titration is not feasible^10–12^. Similarly, B-/plasma-cell therapies currently lack practical levers for patient dose adjustment and rapid reversal, and can be limited by host immune responses due to the use of cell engineering strategies that typically employ viral vectors^13,14^. Across these genetic engineering approaches, host immune responses, and the absence of retrievability or immediate termination present a persistent barrier where on-therapy control is essential.

Encapsulated cell-based *in situ* “biologics factories” are a transformative approach for therapeutic delivery that balances durability with a continuous production and release profile, tunable dosing, and multiplexing^15^. To enable scaling to large patient populations, these biologics factories should be engineered from durable, highly potent allogeneic cell lines (i.e., ARPE-19 cells) and encapsulated in immunoisolating biomaterials to mitigate immune clearance and sustain efficacy^16,17^. Encapsulated cell therapies have demonstrated effectiveness in preclinical models of diabetes, autoimmunity, and oncology, and FDA approval for MacTel type 2 (ENCELTO; Neurotech) sets a first FDA approved precedent^18–21^. However, despite significant advances in immunomodulatory and antifibrotic biomaterials, broad translation remains limited by the foreign body response (FBR), which drives fibrotic matrix deposition that restricts oxygen, nutrient, and therapeutic transport, reducing long-term viability and output^22–26^. Durable function, therefore, requires biomaterials that both mitigate fibrosis and provide sufficient permeability for large biologics (e.g., ∼150 kDa mAbs). Dispersed alginate microcapsules, one of the most common methods for administering encapsulated cells, have shown efficacy but complicate dose control, preclude reliable retrieval, and complicate re-administration, motivating the development of single, implantable, retrievable macrodevices^20,27,28^. Many macrodevice designs, however, have been limited by mass-transport constraints, bulky form factors, and exacerbated foreign body responses^23,29–32^. Addressing these constraints requires an integrated platform that couples an antifibrotic material with a high-productivity cell chassis and a retrievable, mass-transport-optimized macrodevice.

To address these challenges, we developed a modular ‘biologics factory’ by independently optimizing the encapsulating biomaterial, the cellular chassis, and the device design. Using microcapsules as a high-stringency test bed, we first conducted a systematic *in vivo* materials screen, identifying an immunomodulatory alginate that supports year-long serum mAb titers at efficacious levels. Mechanistic interrogation via single-cell RNA-seq of immune cells revealed that this material promotes a locally tempered, pro-resolving niche with reduced inflammation and fibrosis to prevent foreign body responses that typically lead to implant failure. We further demonstrated platform versatility by delivering multiple clinically relevant mAbs from high-potency cell lines developed leveraging cell engineering and synthetic biology approaches. Then, we integrated this lead formulation into a minimally invasive, retrievable macrodevice that enables dose titration, on-demand retrieval, and re-implantation. Finally, we validated the long-term function of our encapsulated cell ‘biologics factory’ platform in non-human primate 6-month durability, dose escalation, and safety studies. To our knowledge, this is the first demonstration of year-long *in vivo* secretion of functional mAbs from an allogeneic, retrievable, and dose-tunable cell-therapy platform. Together, these advances outline a practical route to safe, durable, and tunable cell-based antibody therapy.

## Results

### Modified alginates enable year-long sustained delivery of mAb from encapsulated engineered cells *in vivo*

#### Engineering genetic architectures for high-potency mAb expression from allogeneic clones

To facilitate rapid clinical translation, we chose ARPE19 cells as our cell chassis for generating “biologics factory” for sustained antibody delivery, which are already a component of an FDA approved encapsulated cell therapy (ENCELTO; Neurotech)^33^. ARPE-19 cells are well-suited for allogeneic implantation due to their non-tumorigenic profile, contact-inhibited growth, and established safety record^16,34^. To improve monoclonal antibody (mAb) output (cell potency), we systematically screened genetic architectures by evaluating different co-expression strategies, comparing IRES, P2A, and a dual promoter system to vary heavy- (HC) and light-chain (LC) ratios in stably transfected cells. We found that overexpressing the LC with an IRES linker increased normalized per-cell productivity by ∼3-fold relative to equimolar (P2A) LC/HC expression (*p* < 0.0001; **Fig. 1a**). While dual promoter and IRES systems yielded comparable secretion levels, we selected the IRES system as the optimal construct design due to its smaller genetic footprint. Using this optimized construct architecture, we engineered ARPE-19 cells to produce 3BNC117, a clinically relevant broadly neutralizing antibody against HIV^35,36^ (**Fig. 1b**). By implementing a monoclonal cell isolation and screening pipeline, we identified a stable high-performing clone (ADC26.2) with 3.2-fold higher per cell productivity (pg/cell/day) than the original polyclonal parent population, a gain consistently achieved across multiple other mAb targets (*p* < 0.0001; **Fig. 1c, Extended Fig. 1, Supplementary Fig. 1**). Finally, TZM-bl assays confirmed that ADC26.2-derived 3BNC117 maintained potent neutralization activity against three HIV-1 Env-pseudotyped viruses (**Fig. 1d**).

**Figure 1.**
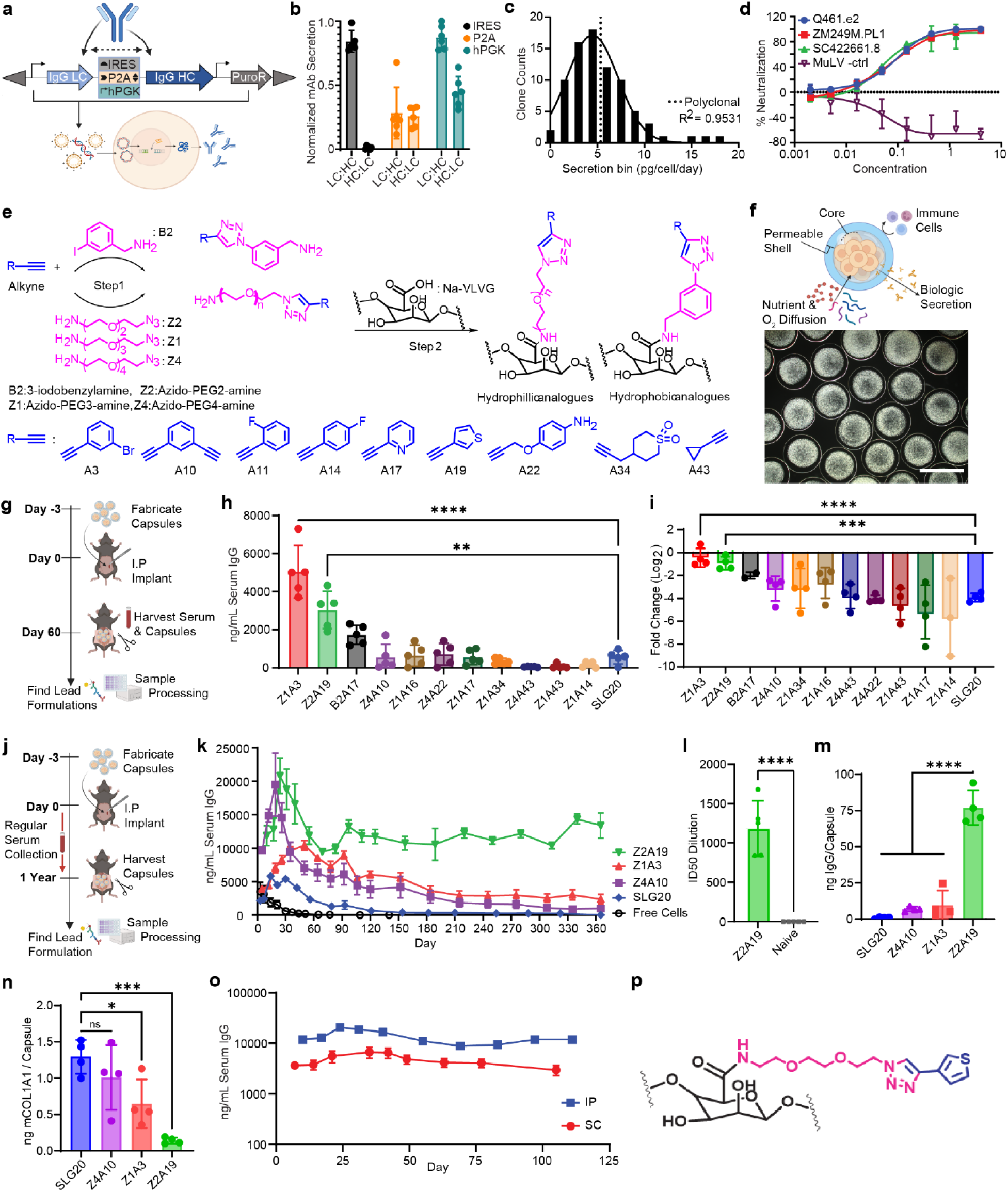
Immunomodulatory alginate enables year-long sustained monoclonal antibody delivery. **a-d,** Cell Line Engineering and Validation: **a,** Schematic of the expression vector design being used to engineer cells. **b,** Six different genetic construct variants for mAb production were tested, and normalized to per cell secretion (±SD, n=7 for group 1, n=8 for groups 2-5). **c,** Histogram of per-cell secretion of clones isolated from an engineered polyclonal population fitted with a Gaussian curve, polyclonal population per cell secretion shown with a dotted line. **d,** Neutralization of Env-pseudotyped viruses in vitro by 3BNC117 in a TZM-bl assay (n=3). **e,** A library of immunomodulatory alginates was created via a two-step chemical synthesis. **f,** Schematic of the core-shell capsules and a 4x dark field transmittance image of capsules containing ARPE-19 cells (Scalebar = 2 mm). **g,** 60-day screening timeline in immunocompetent C57BL/6 mice. **h,** Serum titers at day 60 identify lead formulations (e.g., Z2A19, Z1A3) that sustain antibody delivery (±SD, n=5). **i,** Fold change (log_2_-transformed) in capsule productivity post-explant relative to before implant (±SD, n=4). **J-n,** 1-Year Long-Term Validation: **j,** 1-year timeline for identifying lead material in in B-cell–deficient mice. **k,** Serum 3BNC117 concentrations in B-cell–deficient mice over 1 year (±SEM, All groups n=5, Z4A10 n=4 after day 50). **l,** HIV pseudovirus neutralization (TZM-bl assay) by serum collected at 6 months (±SD, n=5). **m,** Capsule productivity post-explant at day 365 (±SD, n=4). **n,** Collagen overgrowth quantified after one year (±SD, n=4). **o,** Serum 3BNC117 concentrations over 100 days after SC or IP implantation of Z2A19 capsules in B-cell–deficient mice (±SEM, n=5). **p,** Chemical structure of lead triazole-modified alginate, Z2A19. ns (P>0.05), * (P≤0.05), ** (P≤0.01), *** (P≤0.001), **** (P≤0.0001).

#### Screening immunomodulatory alginate for sustained mAb delivery

To identify a lead immunomodulatory biomaterial that supports sustained mAb delivery from encapsulated cells, we designed and synthesized a focused library of eleven small molecule-modified alginate formulations that were previously reported to mitigate fibrosis as part of >200 formulation screen^25,37^. Each formulation was modified with a distinct triazole-containing small molecule via click-chemistry synthesis and amide-coupling conjugation, systematically varying the aromatic head group and PEG linker length to explore hydrophilic and hydrophobic chemical space (**Fig. 1e**). Unlike the prior high-throughput screens that identified antifibrotic alginates using pooled materials screens and insulin-scale payloads, we screened each formulation individually under the dual selective pressure of fibrosis mitigation and permeability to large (∼150 kDa) mAb biologics. Initial *in vitro* validation of our 1.5 mm core-shell (**Fig. 1f**) capsule fabrication process confirmed lot consistency, high cell viability, and functional mAb release (**Supplementary Fig. 2**). To validate baseline *in vivo* durability and dose titratability, we implanted encapsulated cells into immunodeficient (NSG) mice. In this environment, which lacks an effective FBR, the capsules remained stable and showed a linear correlation (R^2^ = 0.9531) between the number of implanted cells and measured serum mAb titers (**Extended Fig. 2a-c**). Applying this workflow to our modified alginate library, all formulations supported >95% viability and efficient mAb diffusion. Notably, Z2A19 achieved the highest release kinetics, matching the performance of unencapsulated cells (*p* = 0.759; **Extended Fig. 3a-b**).

To evaluate long-term therapeutic potential for each formulation, we performed a 60-day head-to-head *in vivo* screen in immunocompetent C57BL/6 mice by implanting encapsulated ADC26.2 cells into the intraperitoneal (IP) space, using the magnitude of terminal serum antibody concentration as the primary metric of success (**Fig. 1g**). After 60-days, the two lead formulations Z1A3 and Z2A19 yielded serum mAb titers that were 5- to 14-fold higher on average than previously identified gold standard formulations, Z1A34^25^ and Z4A10^37^ (**Fig. 1h**). To determine if serum titers correlated with the health and continued productivity of the encapsulated cells, we compared the relative change in mAb productivity of capsules at explantation to pre-implant levels. Capsules fabricated from Z1A3 (*p* < 0.0001 versus SLG20) and Z2A19 (*p* < 0.0001 versus SLG20) exhibited less functional decline, retaining on average 96% and 57% of their pre-implant productivity, respectively, compared to just 6% for SLG20 alginate and 13% (Z1A34) to 19% (Z4A10) for prior leads (**Fig. 1i**). Given the critical role that protein adsorption and exacerbated collagen overgrowth plays in the FBR^38^, the sustained performance of Z2A19 is likely driven by the material’s unique surface chemistry, which exhibited 22-fold lower collagen deposition on explanted capsules (*p* < 0.0001), and significantly reduced albumin adsorption (*p* = 0.0012) compared to SLG20 controls during *in vitro* studies (**Extended Fig. 3c-d**).

Based on the 60-day screen results, we advanced the top two candidates, Z1A3 and Z2A19, to a one-year study compared to the best performing gold standard historical lead, Z4A10 (**Fig. 1j**). To assess long-term material performance independent of anti-drug antibodies (ADAs), we utilized B-cell deficient (BCD) mice, which retain a functional fibrotic response while exhibiting translational mAb pharmacokinetics^39^. While all three modified alginate formulations initially produced high mAb titers, only the Z2A19 group maintained stable serum titers for the entire year (**Fig. 1k**). Titers from the Z2A19 group were 5.6-fold, 13.7-fold, and 52.6-fold higher than those of the Z1A3, Z4A10, and SLG20 groups, respectively. Confirming the longevity of the platform, serum harvested six months post-implantation from Z2A19 mice demonstrated potent HIV neutralization. Treated mice achieved an average ID_50_ of 1183, which was significantly greater than naive serum controls (*p* < 0.0001; **Fig. 1l**), validating the durable secretion of bioactive 3BNC117. Upon explant after 1 year, we observed that the microcapsules remained structurally intact, demonstrating remarkable durability (**Extended Fig. 3e-f**). Furthermore, Z2A19 capsules maintained significantly higher mAb productivity at approximately 75 ng per day per capsule compared to 6-10 ng per day per capsule across other groups (*p* < 0.0001; **Fig. 1m**) and accumulated 4- to 10-fold less surface collagen compared to the other groups (*p* = 0.0005 versus SLG20) at explant **(Fig. 1n)**. Finally, subcutaneous (SC) and intraperitoneal (IP) implants demonstrated long-term durability for over 100 days, highlighted by comparing the final and initial timepoints. At the study’s endpoint, the titer for the SC group was 97% of its initial measured value, while the IP group’s titer was 102% of its initial value **(Fig. 1o)**. This systematic *in vivo* screening thus identified Z2A19 **(Fig. 1p)** as the lead small-molecule for modifying alginate for long-term delivery of mAbs from an encapsulated cell platform across multiple *in vivo* compartments.

#### Sc-RNA-seq reveals that Z2A19 promotes an anti-inflammatory, pro-resolving state

To further investigate the underlying antifibrotic and immunomodulatory mechanisms of our lead formulation, we first examined the composition of cellular overgrowth on explanted Z2A19 and unmodified alginate control capsules (SLG20) via immunofluorescence staining. Both materials exhibited qualitatively comparable recruitment of CD68+ macrophages, which are known to play a central role in the foreign body response and fibrosis. However, there was a decrease in alpha-smooth muscle actin (α-SMA) positive myofibroblasts surrounding Z2A19 compared to SLG20 (**Fig. 2a & Supplementary Fig. 3)**^40,41^. The reduced accumulation of α-SMA+ fibroblasts in Z2A19 indicated attenuation of fibrotic remodeling but did not reveal which immune cell populations or signaling pathways are responsible. We therefore hypothesized that Z2A19 actively alters the local immune landscape, shifting the local immune niche and macrophage polarization toward anti-inflammatory phenotypes that limit fibroblast activation and extracellular matrix (ECM) deposition.

**Figure 2.**
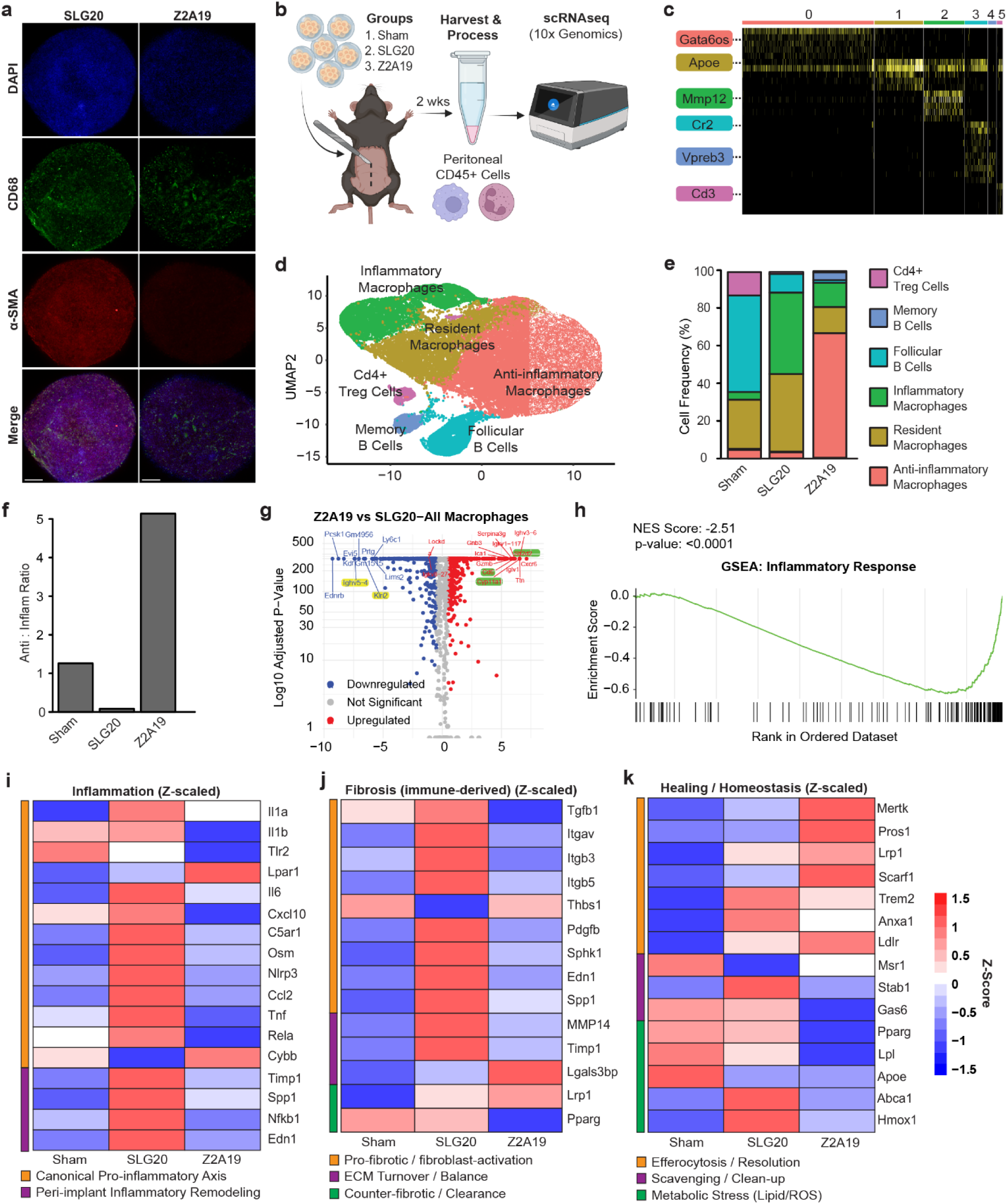
Single cell sequencing analysis in sham mice, mice implanted with Z2A19, and mice implanted with SLG20. **a,** Representative immunofluorescent images of unmodified SLG20 and Z2A19 capsules stained for CD68 (green), DAPI (blue), and A-SMA (red). Scale bars = 200 μm. **b,** A study schematic and timeline for the single cell RNA sequencing study. **c,** Heatmap of the highest expressed genes used to label cell clusters. There are 6 cell clusters. Each row is a different gene, with yellow being more highly expressed and black less expressed. Genes highlighted on the left of graph are representative genes. **d,** UMAP of integrated single-cell RNA sequencing data showing the distribution of 6 clusters. Each dot is a single cell. **e,** Frequency bar graph of anti-inflammatory macrophages across 3 conditions. **f,** Frequency bar graph of anti-inflammatory to inflammatory macrophage ratio across 3 conditions. **g,** Volcano plot as a comparison between mice implanted with Z2A19 versus mice with implanted SLG20 capsules. Red dots are upregulated genes. Blue dots are downregulated genes. Gray dots are genes that are not statistically different between conditions. Green-highlighted genes are those associated with anti-inflammatory properties. Yellow-highlighted genes are those associated with inflammatory properties. **h,** Gene set enrichment analysis for differential genes between mice implanted with Z2A19 versus mice with implanted SLG20 capsules. **I-k,** Heatmap comparison of Z-score of each sample as indicated in the graph relative to the average of the three samples of significant hallmark **i,** inflammation-related genes **j,** fibrotic-related genes **k,** and Healing / Homeostasis related genes organized by function.

To elucidate the mechanisms underlying Z2A19’s antifibrotic properties, we performed single-cell RNA sequencing (scRNA-seq) on CD45+ sorted immune cells harvested from the IP space of mice implanted with either Z2A19 or SLG20 capsules, as well as sham-surgery mice acting as a baseline control (**Fig. 2b**). This approach enables unbiased transcriptional profiling of individual immune cells, allowing discrimination among macrophage, lymphocyte, and stromal populations and identification of pro- versus anti-inflammatory transcriptional signatures. Based on this framework, sequencing analysis revealed a predominantly myeloid-derived population with six clusters (**Supplementary Fig. 4-10**).

To further characterize these cell clusters, we investigated highly expressed genes as well as canonical markers (**Fig. 2c**). The combination of these markers revealed an anti-inflammatory -like macrophage population, a resident macrophage population, an inflammatory -like macrophage population, a follicular B cell population, a memory B cell population, and a CD4 T cell population (**Fig. 2d**). After identifying these six cell clusters, we next quantified their relative abundance across the three experimental conditions: sham mice, mice implanted with Z2A19, and mice implanted with SLG20 alginate (**Fig. 2e**). In the sham mice, follicular B cells were the most common type of cells followed by resident macrophages, and then T cells (**Fig. 2e**). In the mice implanted with SLG20, there was a shift from B cells to predominantly inflammatory macrophages (**Fig. 2e**). In mice implanted with Z2A19, the immune landscape shifted from a B cell–dominant profile to one enriched in anti-inflammatory-like macrophages (**Fig. 2e**). Moreover, the anti-inflammatory:pro-inflammatory macrophage ratio was higher in mice implanted with Z2A19 (5.13) than SLG20 controls (0.083) resulting in a more than a 60-fold difference (**Fig. 2f**). These findings indicate a less inflammatory immune environment potentially arising from reduced pro-inflammatory recruitment and/or enhanced anti-inflammatory polarization.

Given that the Z2A19 appeared to promote a more anti-inflammatory state, we next investigated gene expression profiles of macrophages between the Z2A19 and SLG20 capsule-treated mice (**Fig. 2g**). Analysis of macrophage subsets showed numerous up-regulated genes associated with anti-inflammatory states (Cd6, Cyp11a1, Sh2d2a) and down-regulated genes associated with inflammatory states (Klri2, Ighv5-4). The change in cellular populations, coupled with the expression of pro- and anti-inflammatory genes, led us to investigate pathway dynamics further via gene set enrichment analysis (GSEA) of inflammatory genes (**Fig. 2h**). Compared with SLG20 controls, macrophages from Z2A19–implanted mice displayed a broad, global downregulation of inflammatory genes. GSEA of inflammatory genes showed an NES score of −2.51 with a p-value of <0.0001, indicating that many inflammatory genes were downregulated.

Lastly, given the gene-set analysis differences between Z2A19 and SLG20 alginate, we investigated individual genes in the inflammatory set, finding stark changes in gene expression in Z2A19 implanted mice compared with SLG20 implanted controls (**Fig. 2i-k**). For example, many inflammatory genes that were upregulated with SLG20 were downregulated with Z2A19, including Il1b, TLR2, CXCL10, Tnf, Rela, Ccl2, and Timp1 (**Fig. 2i**). These data are consistent with a lower inflammatory transcriptional profile in Z2A19 relative to SLG20. In addition, a more comprehensive GSEA analysis comparing Z2A19 relative to SLG20 for all hallmark gene sets, was performed showing a stark trend towards inhibition of pro-inflammatory immune cell signaling pathways such as TNF-α, IL6, IL2, TGF-ϐ, and IFN-γ all with a p-value <0.002 (**Extended Fig. 4**). Epithelial mesenchymal transition was also significantly down regulated (p-value < 0.0001), which is important in the initial phases of wound healing, but chronic activation is indicative of scarring and fibrosis^42^. Using curated fibrosis and healing/homeostasis gene sets^23,43,44^, we observe a clear shift with the downregulation of pro-fibrotic, fibroblast-activating signals (e.g., TGF-ϐ1, THBS1, SPP1) and upregulation of pro-resolving/homeostatic markers (e.g., MERTK, AXL) (**Fig. 2j, k**). Together, these results point towards Z2A19 promoting an anti-inflammatory local niche around the capsules, driving tissue homeostasis and reduced fibrotic overgrowth when compared to SLG20, whose immune profile is more consistent with chronic inflammation driving exacerbated foreign body response, ultimately leading to thick collagen-dense fibrotic overgrowth on the SLG20 capsules and resulting in cell death and reduced mAb secretion over time. This is consistent with our experimental results showing reduced collagen build-up, lower α-SMA+ signal, increased cell survival, mAb secretion, and durable titer levels over time from Z2A19 capsules relative to SLG20, linking the local immune state to reduced fibrotic remodeling^41,45,46^.

#### Modified alginates can support the delivery of a range of bioactive mAbs

There are a large number of disease indications with effective mAb treatments that are FDA-approved or in clinical development, that would benefit from this cell-factory platform^47^. Many FDA approved mAb sequences differ only in the variable regions of the light and heavy chains, which function to bind the antigen of interest, meaning that our optimized cell engineering strategy is likely to port over to other mAbs (**Fig. 3a**). This led us to implement a standardized pipeline for the development of several clinically relevant mAb producing cell lines as a demonstration of platform versatility (**Fig. 3b**). We were able to engineer ARPEs to stably produce 13 different mAbs across oncology, infectious disease, and autoimmune disease indication to name a few (**Fig. 3c**). Serum-free media supernatant harvested from select antibody-producing cell lines was used to verify secretion of the correct mAb sequence from each cell line using mass spectrometry (**Fig. 3d**), with QC and abundance summaries reported in **Supplementary Fig. 11**. This analysis also allowed us to assess the complete secretome from our cell lines including a naïve ARPE-19 control and analyze the top constituents thereof (**Fig. 3e**). It was shown that the mAb of interest was generally the most highly secreted protein and in the case of the PGT121 expressing clone, it was ∼34 fold more highly abundant than the next highest protein. This, combined with the observation that the top secretome constituents were generally associated with ECM homeostasis, indicates likely minimal off-target effects from other produced proteins (**Supplementary Table 2**). To demonstrate that the produced mAb is correctly folded and functional, binding of the produced antibody to the antigen of interest was shown either via TZM.bl assay or via ELISA (**Fig. 3f-j**). All tested cell lines were shown to secrete functional mAbs, and the anti-HIV mAbs 3BNC117 and PGT121 were bioactive, neutralized pseudovirus, preventing cell infection, similarly to the respective clinical-grade control antibody. To demonstrate that dosing could be precisely controlled, the number of capsules administered per group was adjusted based on each cell line’s specific productivity (pg/cell/day) to normalize the total daily mAb dosage (ng/day) across all experimental cohorts (**Fig. 3k**).

**Figure 3.**
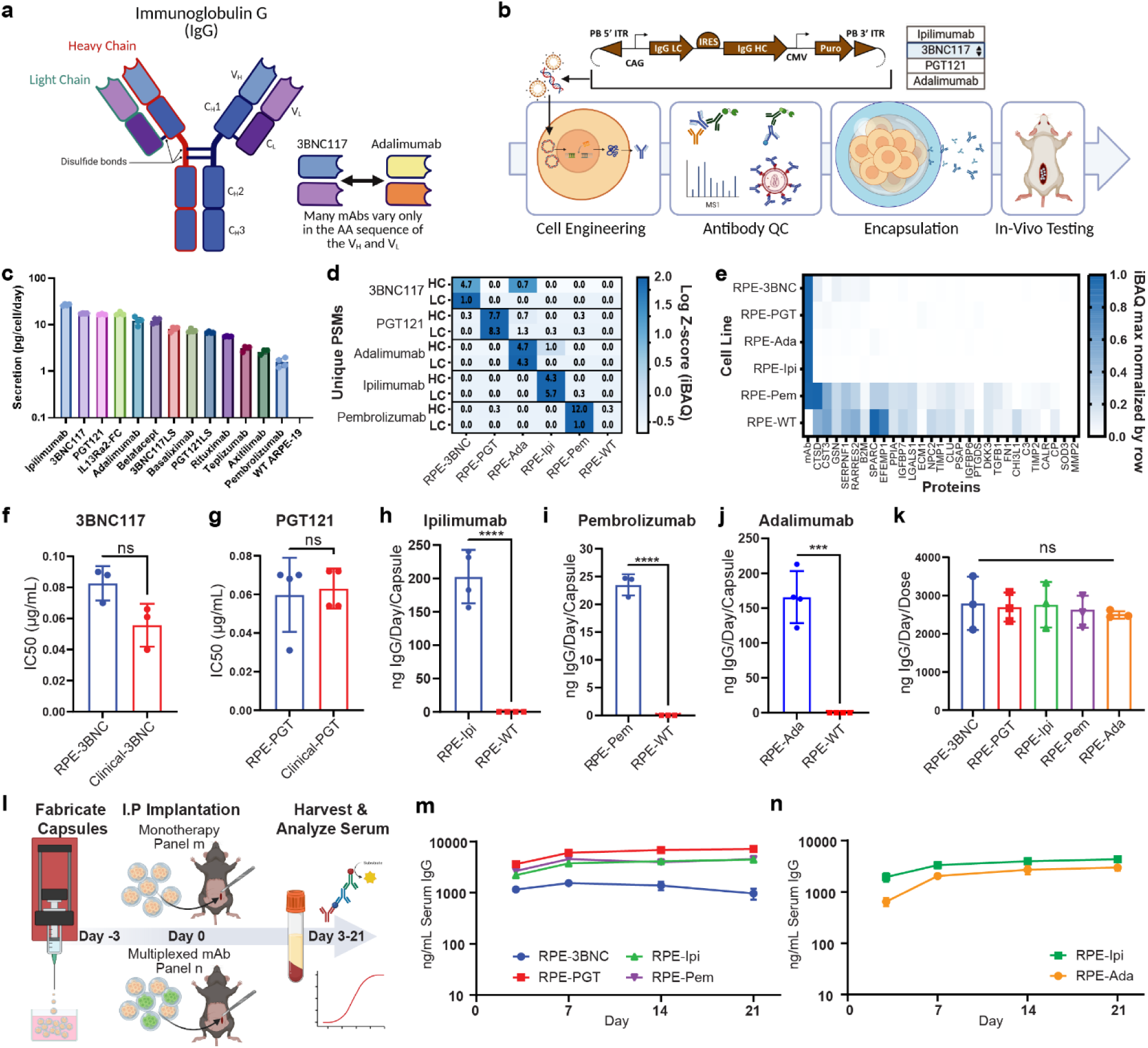
Therapeutic cells encapsulated in Z2A19 remain efficacious for a broad range of mAbs. **a,** General structure of an IgG monoclonal antibody. Therapeutic diversity is achieved by modifying the amino acid sequences of the heavy and light chain variable regions. **b**, General workflow describing the ability of Engineered ARPE-19 cells to be engineered to produce a range of mAbs followed by quality control (QC), encapsulation, and *in vivo* testing. **c**, Per cell productivity data of mAbs and FC-fusion proteins produced by engineered ARPE-19 cells (n=4). **d**, Mass spectrometry confirms the identity and high relative abundance of each specific mAb via unique peptide spectrum matches, annotated in each box, and by statistical significance indicated by Log(z-score) of IBAQ score across groups and box shading (n=3). **e**, Proteomic analysis of the secretome from each cell line. Each row is max normalized by IBAQ score (n=3). **f-g**, TZM-bl HIV pseudovirus neutralization assays using supernatant from encapsulated ARPE-19 cells engineered to secrete **f**, 3BNC117 (±SD, n=3) or **g**, PGT121 (±SD, n=4). **h-j**, Antigenicity ELISA quantification of antibody secretion from encapsulated ARPE-19 cells engineered to produce **h**, ipilimumab (±SD, n=4), **i**, pembrolizumab (±SD, n=3), or **j**, adalimumab (±SD, n=4). **k**, Pre-implant total productivity per planned dose for each mAb (n=3). The number of capsules per dose was adjusted to normalize total output to a target of ∼2.5 μg/day before implantation. **l**, A schematic illustrating single mAb and multiplexed mAb delivery *in vivo* from Z2A19 encapsulated cells. **m**, Serum IgG concentrations over 3 weeks in C57BL/6 mice implanted with the productivity-matched doses shown in k (±SEM, n=4 for 3BNC117, n=5 for all other groups). **n**, Serum IgG concentrations over 3 weeks in RAG2 mice from a single multiplexed implant co-delivering ipilimumab and adalimumab at matched pre-implant productivity (±SEM, n=5). ns (P>0.05), *\*\*\*\* (P≤0.0001)*.

With this normalized dosing strategy, we next evaluated whether Z2A19 could serve as a modular platform for diverse therapeutic needs by testing its ability to support both monotherapy and multiplexed mAb delivery (**Fig. 3l**). Monotherapeutic delivery of four distinct mAbs resulted in sustained serum levels across all cohorts (**Fig. 3m**). We further extended this approach to combination delivery by co-implanting capsules producing ipilimumab and adalimumab in RAG2-/- mice, a model chosen to avoid strain-specific clearance of adalimumab (**Supplementary Fig. 12**). Both antibodies were successfully quantified via specific ELISAs, showing stable co-delivery and comparable pharmacokinetics with titers exceeding 1 µg/mL (**Fig. 3n**).

#### Macrodevice integration enables minimally invasive, retrievable cell delivery

A significant barrier to the clinical translation of encapsulated cell therapies, particularly microcapsules, is the difficulty in retrieving or replacing implants^28^. The ability to retrieve implants is critical for managing the safety of allogeneic engineered cell therapies, as it allows for the mitigation of adverse side effects and provides greater control over dosing and exposure duration^48,49^. To address this challenge, we integrated Z2A19 alginate-encapsulated cells into a novel, retrievable macrodevice and evaluated *in vivo* performance (**Fig. 4**). The macrodevice is composed of a 3D printed hexagonal lattice that provides mechanical stability for a hydrogel core while enabling minimally invasive trocar injection and retrieval (**Fig. 4a, Extended Fig. 5a-d**). Its ≥1 mm pores are designed to permit the diffusion of waste, nutrients, and therapeutics while minimizing tissue overgrowth and scar formation^50^. *In vitro*, 3BNC117-producing ARPE-19 cells exhibited equivalent viability (≥96%) and per-cell productivity when loaded as a bulk alginate slab or as 1.5-mm capsules within the macrodevice (**Fig. 4b; Extended Fig. 5d–e**). However, *in vivo*, the capsule-loaded configuration produced higher sustained titers than the slab-loaded configuration (**Extended Fig. 5f**), providing a rationale for capsule loading of macrodevices in subsequent studies.

**Figure 4.**
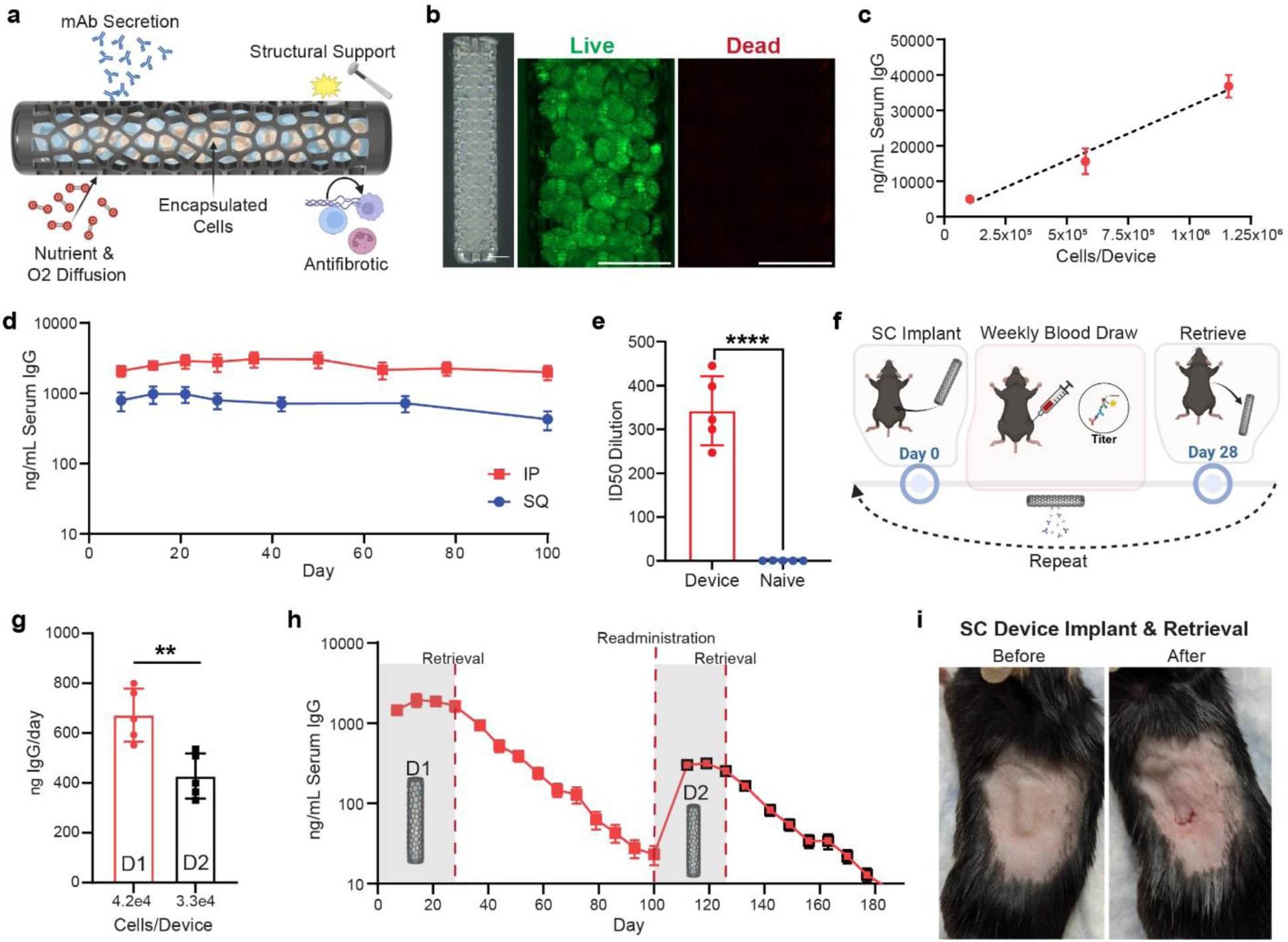
A retrievable macrodevice enables long-term, tunable, and effective in vivo antibody delivery. **a,** Schematic illustrating the device design, showing an immunomodulatory alginate core containing encapsulated cells within a retrievable lattice structure. **b,** Representative live (green) / dead (red) staining of capsule-loaded macrodevice-encapsulated cells 3 days post-loading (scale bar = 2 mm). **c,** Dose-dependent antibody delivery from macrodevices encapsulated with 1×10⁷ cells/mL (n=5), 4×10⁷ cells/mL (n=4), or 8×10⁷ cells/mL (n=5), with serum 3BNC117 concentrations (±SEM) correlating to the number of implanted cells at day 28 (R^2^=0.865). **d,** Serum 3BNC117 concentrations following subcutaneous (SC) or intraperitoneal (IP) implantation in B-cell–deficient mice over 100 days (±SEM, n=5). **e,** Neutralization of HIV pseudovirus in TZM-bl assay by serum collected at day 28 from macrodevice-implanted mice compared with naïve controls (±SD, n=5). **f,** Schematic of SC implant, blood collection, retrieval, and re-implantation study design. **g,** 3BNC117 production rates measured prior to implantation for the initial macrodevice (D1) and replacement macrodevice (D2). ±SD, n=5. **h,** Serum 3BNC117 concentrations in the retrieval/re-implantation cohort; D1 retrieved at day 28, D2 re-implanted at day 100 (±SEM, n=5). **i,** Representative images of SC implant site before and after macrodevice retrieval. ** (P≤0.01), **** (P≤0.0001).

Dosing of macrodevices can be controlled by varying the density of encapsulated cells (cells/mL), adjusting the total implant volume by changing macrodevice length, or implanting multiple devices. To demonstrate tunable dosing, we implanted macrodevices of identical size and volume with varying densities of encapsulated 3BNC117-producing cells (1-8×10⁷cells/mL) into the IP compartment of BCD mice. After 28 days, serum mAb concentrations showed a consistent dose-dependent relationship (R^2^=0.865) with the number of encapsulated cells per macrodevice, resulting in serum concentrations ranging from 5-48 µg/mL (**Fig. 4c, Extended Fig. 6a-c**). Representative 100-day explants from the titration study demonstrate device retrievability and protection of capsules housed within, evident by the lack of cellular and fibrotic overgrowth on the capsule surfaces (**Extended Fig. 6d**).

To assess the long-term *in vivo* durability of the macrodevices, we implanted 3BNC117-producing macrodevices either subcutaneously (SC) or intraperitoneally (IP) in BCD mice, tracking serum concentrations for 100 days. Sustained antibody levels were achieved at both implantation sites for 100 days, with IP implants showing remarkable stability (96% of initial value) and SC implants maintaining a durable, albeit lower, titer (54% of initial value) (**Fig. 4d**). Supplementary data shows that implants from the IP cohort maintained high titers through one year (**Extended Fig. 6e**). Furthermore, secreted antibody function was confirmed with serum collected from the IP group one-month post-implantation, showing potent HIV neutralization in a TZM-bl pseudoviral assay with an average ID_50_ of 342, contrasting the negligible activity (average ID_50_ of 0.2) in serum from naïve mice (*p* < 0.0001; **Fig. 4e**).

A key advantage of this platform is the ability to terminate or modulate therapy on demand. To demonstrate that therapy can be precisely controlled, we performed an implant, retrieval, and re-administration study (**Fig. 4f-i**). To demonstrate retrieval and therapeutic termination, macrodevices were implanted SC and then retrieved after 28 days (**Fig. 4f**). Then, to further highlight the platform’s ability to modulate dosage, we re-implanted the same mice with a second macrodevice engineered to produce less mAb (**Fig. 4g**). Retrieval of the first device after 28 days prompted an immediate shift from steady state pharmacokinetics to a terminal clearance phase, with titers declining from an initial concentration of 1,445 ng/mL to 23 ng/mL by 60 days post-retrieval (**Fig. 4h**). Re-implantation of the second macrodevice engineered to produce less mAb resulted in a lower therapeutic level, measuring 301 ng/mL compared to the initial implant’s 1,445 ng/mL (**Fig. 4h**). Finally, images taken before and after the procedure illustrate that the SC macrodevices can be retrieved via a minimally invasive method that does not require sutures for closure (**Fig. 4i**).

#### Modified alginates and macrodevices are biocompatible and well-tolerated in healthy mice

To assess the safety of the biomaterial portion of our delivery platforms, we implanted acellular Z2A19 modified alginate capsules and macrodevices subcutaneously in healthy C57BL/6 mice and monitored them for one month (**Fig. 5**). In this assessment, we evaluated systemic health via changes in body weight and analysis of blood chemistry, and evaluated the local tissue response using histology of the implant site.

**Figure 5.**
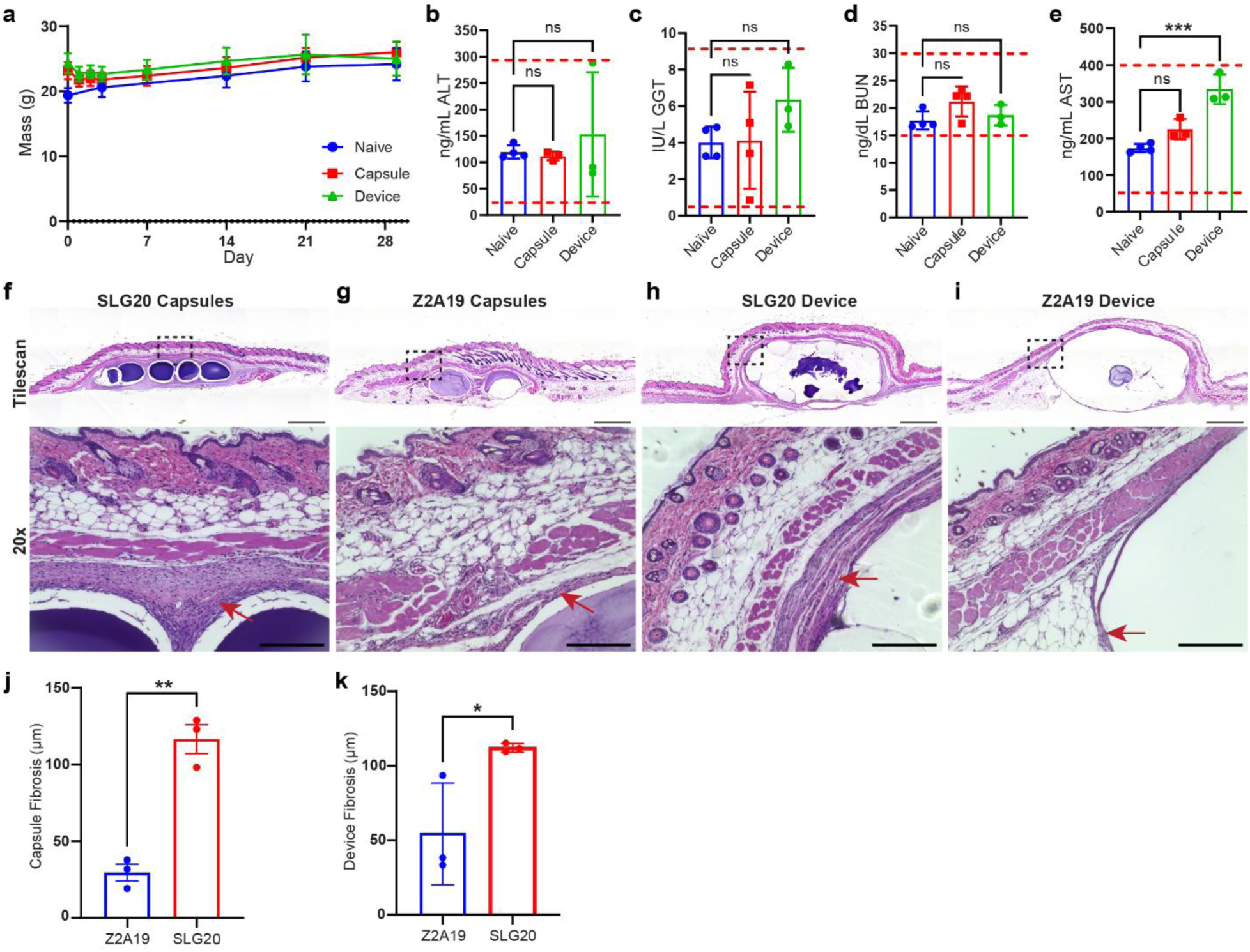
Z2A19 capsules & macrodevices are well tolerated in healthy mice. Biocompatibility was evaluated one month after subcutaneous implantation of acellular Z2A19 capsules or macrodevices, compared with naïve controls. **a,** Body weight was monitored weekly (±SEM, n=4 naïve and capsule, n=3 device). **b-e,** Serum chemistry at day 28 showing **b,** alanine aminotransferase (n=4 naïve, n=3 capsule and device, ALT) **c,** gamma-glutamyl transferase (n=4 naïve and capsule, n=3 device, GGT) **d,** blood urea nitrogen (n=4 naïve and capsule, n=3 device, BUN), and **e,** aspartate aminotransferase (n=4 naïve, n=3 capsule and device, AST). Red dotted lines denote reference physiological ranges. Error bars = ±SD. **f-i,** Representative hematoxylin and eosin (H&E)–stained sections of subcutaneous tissue. Top: tile scans; bottom: higher-magnification (20x) regions corresponding to boxed areas. **f,** SLG20 capsules. **g,** Z2A19 capsules. **h,** SLG20 macrodevice. **i,** Z2A19 macrodevice. Scale bars: 1 mm (tile scans), 200 μm (20x). **j,** Quantification of fibrotic capsule thickness of Z2A19 versus SLG20 capsule implants (±SEM, n=3). **k,** Quantification of fibrotic capsule thickness of Z2A19 versus SLG20 macrodevice implants (±SEM, n=3). Red arrows indicate fibrotic capsule. ns (P>0.05), * (P≤0.05), ** (P≤0.01), *** (P≤0.001).

As a general indicator of health, body weight was monitored throughout the study. All groups, including capsule- and device-implanted mice, exhibited a steady weight gain comparable to naïve (un-implanted) controls with no significant difference at the study endpoint (p=0.4367), suggesting no systemic toxicity (**Fig. 5a**). Blood was analyzed after 1 month for key markers of organ health. While minor variations between groups were noted, such as slightly higher Aspartate Aminotransferase (AST) levels in device-implanted mice compared to naïve (333 vs 173 ng/mL), the levels of Alanine Aminotransferase (ALT), Gamma-Glutamyl Transferase (GGT), blood urea nitrogen (BUN), and AST remained within normal physiological ranges for all of the tested mice (**Fig. 5b-e**). This indicates the implants did not adversely affect liver, kidney, or heart health.

Finally, local tissue biocompatibility at the implant site was confirmed via H&E staining. Both the capsule and macrodevice implants were surrounded by a thin, well-organized fibrous capsule, without evidence of chronic inflammation, necrosis, or extensive immune cell infiltration when compared to tissue from a naïve animal (**Fig. 5f-i & Supplementary Fig. 13**), confirming the implants are well-tolerated locally. Importantly, direct comparison of antifibrotic Z2A19-modified implants with unmodified SLG20 controls revealed a significant difference in fibrotic capsule formation for both capsules (*p* = 0.0013) and devices (*p* = 0.0422). To quantify this response, fibrotic capsule thickness was measured at multiple points per section and averaged across replicates. Z2A19 capsules exhibited fibrotic capsules less than 40 µm thick on average, whereas SLG20 capsules developed capsules around 120 µm (**Fig. 5j**). Similarly, Z2A19 macrodevices were surrounded by capsules averaging less than 60 µm in thickness compared to greater than 110 µm for SLG20 macrodevices (**Fig. 5k**).

#### Platform validation in non-human primates

We evaluated the pharmacology and durability of the Z2A19 formulation in a non-human primate (NHP) via a subcutaneous (SC) implant (**Fig. 6a**). To provide a rigorous “stress test” for the platform’s antifibrotic properties, we utilized cells engineered to produce ipilimumab. As a checkpoint inhibitor, ipilimumab promotes T-cell activation, which creates a more challenging pro-inflammatory environment for the encapsulated cells. A 2 mL SC dose of Z2A19 capsules loaded with ∼4.8×10⁷ total cells was administered following *in vitro* validation of capsule lot productivity and viability (**Extended Fig. 7**). Serum ipilimumab levels peaked at ∼7 µg/mL by day 7, underwent a transient dip around days 60–80, and subsequently recovered to ∼1 µg/mL through day 186 (**Fig. 6b**). This demonstrates over 6 months of continuous drug production from a single administration, well beyond the point at which a simulated equivalent-dose I.V. bolus decayed to sub-nanogram serum levels. To evaluate dose control, we conducted a dose escalation study in two additional primates receiving 6 mL and 12 mL implants, respectively (**Fig. 6c**). Serum ipilimumab concentrations measured 3 days post-implant across all three dose levels (2, 6, and 12 mL) correlated linearly with pre-implant capsule productivity (R² = 0.979), indicating that systemic drug exposure can be predictably tuned by adjusting implant volume and the number of implanted encapsulated cells.

**Figure 6.**
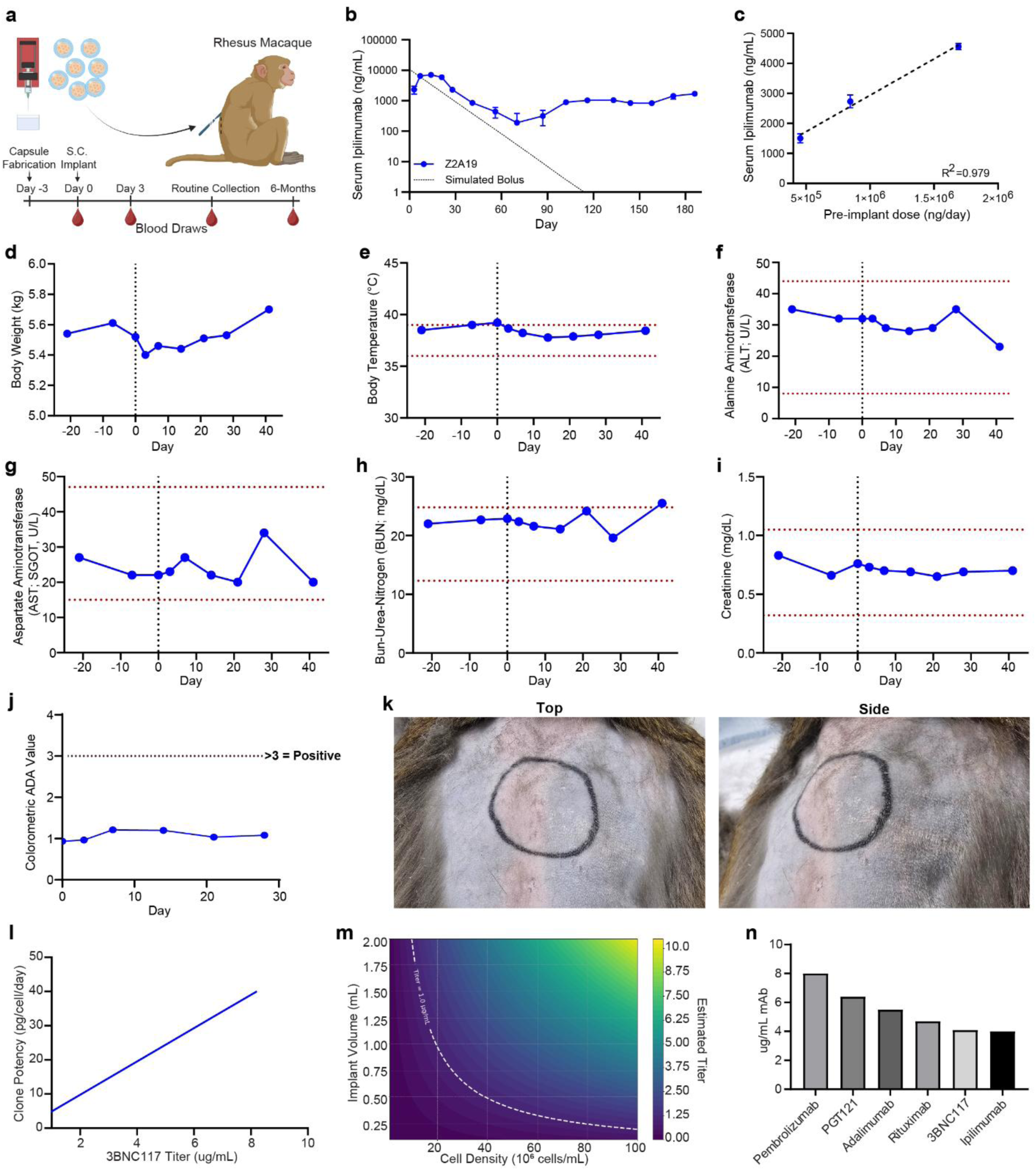
Sustained Z2A19 function and tolerability in non-human primate. **a,** Schematic of non-human primate subcutaneous implant study and timeline (n=1). **b,** Normalized serum ipilimumab concentration (ng/mL) in the NHP (n=1) over time compared to a simulated 0.6 mg/kg I.V. ipilimumab bolus dose (dashed line). **c,** *Dose-dependent serum ipilimumab concentration in NHPs (n=1 per dose) 3 days post-implant as a function of pre-implant daily productivity across three implant volumes (2 mL, 6 mL, and 12 mL) of 1.5 mm Z2A19 capsules encapsulated at 4×10⁷ cells/mL (R²=0.979). Error bars = ±SD from n=3 technical replicates.* **d-i,** NHP general toxicity and blood chemistry data between days −25 and 41 pre-and post-explant. Vertical black dashed lines mark the date of implant and horizontal red dashed lines mark normal reference ranges. **d,** NHP body weight. **e,** NHP body temperature. **f,** Alanine aminotransferase (ALT) levels. **g,** Aspartate aminotransferase levels (AST). **h,** Blood-urea-nitrogen levels (BUN). **i,** Creatinine levels. **j,** Anti-drug antibody levels against Ipilimumab antibodies measured in harvested NHP serum. **k**, Images of the subcutaneous NHP capsule implant site. **l,** Estimated 3BNC117 steady-state titer versus clone potency at a fixed cell density of 4×10⁷ cells/mL and a volume of 2 mL. **m,** Effect of manipulating device volume and encapsulated cell density at a fixed per-cell productivity of 20 pg/cell/day on the expected 3BNC117 titer in humans. **n,** Estimated steady-state titer for multiple mAbs at a fixed volume of 2 mL, cell density of 4×10⁷ cells/mL, and productivity of 20 pg/cell/day.

Blood chemistry and complete blood counts (CBCs) were tracked from days −25 to 41 to assess systemic toxicity and immune activation. The NHP exhibited healthy recovery with stable body temperatures and consistent weight gain post-implant (**Fig. 6d-e**). Monitoring of liver (AST and ALT), and kidney (Creatinine and BUN) markers remained within normal ranges, indicating no organ-specific toxicities (**Fig. 6f-i**). Finally, while CBCs showed initial transient peaks in immune cells counts between days 7 and 21, these levels normalized between days 28 and 41, indicating the resolution of acute post-implant inflammation (**Extended Fig. 7**). Serum quantification of anti-ipilimumab antibodies over the first month post-implant revealed no measurable anti-drug antibodies (ADAs) in the NHP’s serum, further indicating a favorable immune response to the living pharmacy implant (**Fig. 6j**). Finally, images of the NHP’s skin overlying the implant site, two months post-implant, show no visible difference from surrounding unmanipulated tissue, indicating local tolerance over longer periods (**Fig. 6k**)

To contextualize these data for translation, we modeled best-case human steady-state titers using murine- and NHP-informed implant feasibility parameters combined with human-specific clearance rates taken from the FDA label or literature sources (**Supplementary Table 3**)^51,52^. To estimate human titers under best-case conditions, we predicted steady-state serum concentrations assuming zero-order production from the implant and first-order systemic clearance. At a reference setting of 4×10⁷ cells/mL and 2 mL total implant volume, the projected titer for 3BNC117 increases linearly with PCD (**Fig. 6l**). Estimates project ∼4.0 μg/mL at 20 PCD and ∼8.0 μg/mL at ∼40 PCD. Alternatively, with cellular productivity fixed at 20 PCD, iso-titer contours for 3BNC117 visualize operating regimes as a function of implant volume (0.2–2 mL) and encapsulated cell density (1-10×10⁷cells/mL) (**Fig. 6m**). Finally, projections across a panel of clinical mAbs showed steady-state concentrations ranging from ∼4 μg/mL (ipilimumab) to ∼8 μg/mL (pembrolizumab), demonstrating that, while the platform is versatile, the final therapeutic window is ultimately dictated by the specific antibody’s human clearance rate (**Fig. 6n**).

## Discussion

Conventional biologic drug delivery paradigms rely on relatively frequent dosing by injection, characterized by “peak and trough” pharmacokinetics of drug concentration in the body. This results in potential overdosing at the beginning of a drug regimen and underdosing prior to the patient’s next dose, increasing the likelihood of off-target effects and mismanagement of disease at the end of each dosing cycle^2,3,53^. This work establishes a continuous release paradigm, achieving stable, steady-state drug concentrations through a retrievable, dose-tunable, encapsulated cell therapy platform that facilitates stable, long-term delivery of clinically relevant monoclonal antibodies by mitigating the host foreign body response to the implant. This platform leverages parallel advances in cell engineering, biomaterial optimization, and device design to systematically address the critical challenges of functional mAb delivery, foreign body response, and clinical translatability, enabling year-long mAb dose maintenance in mice at clinically relevant levels and sustained mAb release for over 6 months in a non-human primate from a single administration.

As previously outlined, existing strategies for long-acting mAb delivery, such as polymeric depots, pump devices, autologous cell therapies, and gene therapies each face trade-offs between durability, dose control, reversibility, and cost ^5,7,10,12–14,54–57^. In contrast, our cell-based ‘biologics factory’ approach for long-acting mAb delivery overcomes many of these challenges through parallel engineering advances in cell line development, material optimization, and device design. A modular pipeline for engineering high-potency allogeneic clones validated across a range of mAbs whose secretomes are dominated by the biologic of interest (up to ∼34 fold more abundant than the next highest constituent) has been demonstrated, addressing the high cost of autologous cell therapies by utilizing allogeneic cells, highlighting the versatility of the platform, and enabling the possibility of clinically efficacious dosing with reasonable implant volumes. The platform demonstrates a high safety profile and biocompatibility in rodent models and utilizes a cell line that has been translated to the clinic, is generally considered non-tumorigenic, and is contained within a modified hydrogel that does not undergo remodeling by the host, further increasing the safety of the platform ^16,58^. In addition, we engineered a lattice macro-device, to contain the cell-laden hydrogel that is: compatible with minimally invasive outpatient trocar implantation; is designed with ≥1 mm lattice pores associated with reduced granuloma bridging and foreign body scarring ^50,59^; allows for facile dose tuning that is linearly correlated with contained cell number (R² = 0.865); and enables on-demand modulation or cessation, without genetic safety switches or invasive surgery, by minimally invasive retrieval and subsequent reimplantation. Interestingly, macrodevice contained capsules explanted at 100 days showed minimal cellular attachment and overgrowth (**Extended Fig. 6d**), and both SC and IP implants sustained stable titers over this period. Combined with the year-long stable IP titer (**Extended Fig. 6e**), these data suggest that the lattice design modulates cellular infiltration, while preserving the performance of the original dispersed microcapsule form factor, warranting further investigation.

While the recent FDA approval of ENCELTO^TM^ validates the feasibility of long-term protein secretion from an autologous engineered RPE cell line *in vivo*^21^, its application is restricted to the immune-privileged ocular compartment and does not address the transport barriers of systemic delivery^60^. Expanding this paradigm to extra-ocular sites requires a fundamental shift in how we identify and validate immunomodulatory materials. We built upon prior large-scale screens identifying antifibrotic modifications for immunocompetent sites but implemented selection constraints essential for mAb delivery. While prior work emphasized insulin-scale payloads and pooled materials for high-throughput searches, they risked bystander effects and crosstalk. We screened formulations individually to prioritize the efficient and sustained diffusion of mAb-scale proteins within their ultimate therapeutic context. This strategy enabled deeper analysis of material performance and FBR mitigation resulting in the identification of Z2A19 a lead material distinct from historical benchmarks that achieved significantly superior dose durability, cell viability, and compatibility with mAb delivery.

The FBR typically begins with protein adsorption and inflammatory recruitment, progressing to macrophage fusion, fibroblast activation, and fibrotic encapsulation ^23,61^, single-cell RNA sequencing reveals that Z2A19 actively diverts this trajectory. When compared to the literature standard SLG20 alginate, it enriches anti-inflammatory, pro-regenerative macrophages and suppresses fibrosis-linked genes (e.g., IL-1b, CCL2). Accordingly, Z2A19 implants display fewer α-SMA⁺ myofibroblasts, thinner fibrotic capsules, less collagen, and superior long-term productivity, linking material chemistry to *in vivo* immunobiology. This approach of engineering an intrinsically immunomodulatory material offers distinct advantages over other strategies, such as co-encapsulating immunosuppressive drugs like dexamethasone that elute over time. These co-encapsulation strategies can reduce fibrosis ^40,62^, but they risk long-term side effects like increased infection susceptibility and fail to address the inherent material limitations of SLG20 alginate, namely the poor diffusion of large biologics that our screening was explicitly designed to overcome. To our knowledge, this is the first biomaterial-based platform to achieve stable therapeutic mAb titers for one year in a mouse model with an intact foreign body response.

The NHP study is a promising indicator of the platform’s clinical translatability, particularly when viewed against the treatment burden imposed by current mAb delivery regimens. For example, Ipilimumab is administered via IV infusion every 3 weeks during induction, with each session requiring 30–90 minutes of chair time at a dedicated infusion center, followed in some indications by maintenance dosing every 12 weeks for up to 3 years. Each visit entails travel, pre-treatment laboratory work, infusion, and post-infusion monitoring, a significant cumulative burden on patients that contributes to treatment non-adherence in chronic disease settings. Moreover, these bolus regimens produce the peak–trough pharmacokinetics that increase risks of off-target toxicity at peak concentrations and diminished efficacy approaching each subsequent dose. By maintaining stable systemic ipilimumab titers (∼1 µg/mL) for over six months from a single implant, the NHP study results presented in this manuscript establish pre-clinical feasibility for a potentially once-yearly treatment paradigm. Notably, this durability was achieved while delivering a pro-inflammatory checkpoint inhibitor that actively challenges the antifibrotic properties of the implant^63^, and predictable dose control was confirmed across a three-dose escalation (R² = 0.979). Such an approach could reduce dozens of infusion center visits to a single outpatient procedure, eliminate the pharmacokinetic volatility inherent to bolus dosing, and meaningfully improve treatment adherence and patient quality of life.

To remain within subcutaneously feasible volumes (≤2 mL), and because oxygen and nutrient availability impose an upper bound on cell density (4×10⁷ cells/mL in our studies), future engineering efforts should prioritize technologies that support higher cell densities (e.g., enhanced oxygenation) or further increases in per-cell productivity to compensate for mAbs with higher clearance rates or exposure requirements^24,26^. In parallel, improvements to the effective potency of mAbs through protein engineering could significantly lower the serum concentration required for efficacy. Although we did not evaluate next-generation protein engineering in this work, current advances in AI-guided mAb engineering motivate parallel efforts in protein and cell engineering alongside device optimization^64^. These bioengineering approaches could be coupled with on-demand, small-molecule-activatable genetic circuits or sense-and-response systems, further enhancing the tunability of this platform^15,65^.

In conclusion, we demonstrate that an integrated platform combining immunomodulatory alginate, high-potency allogeneic cell engineering, and a retrievable macrodevice can achieve year-long, steady-state delivery of therapeutic mAbs in mice and sustained systemic drug levels for over six months in a non-human primate. The platform’s modularity was validated *in vivo* across five clinically relevant mAbs, demonstrating rapid adaptation to diverse therapeutic targets. Key next steps include platform demonstration in disease models, longitudinal immune profiling to evaluate the durability of the anti-inflammatory niche in greater depth, and formal biocompatibility and toxicology assessments required for IND-enabling studies. This approach has the potential to shift the chronic disease management paradigm from frequent injections toward single-administration therapies that can improve outcomes and expand access worldwide.

## Materials & Methods

### Modified alginate synthesis and characterization

#### Modification of VLVG with small molecules

Ultrapure very low viscosity alginate (UP-VLVG) from Pronova was used for modification. A 1 % w/v solution was prepared by dissolving 1.5 grams of UP-VLVG in 150 mL of deionized water. To this solution, a corresponding small molecule (1 equivalent) dissolved in 50 mL of acetonitrile was added dropwise. This was followed by the dropwise addition of 4-(4,6-dimethoxy-1,3,5-triazin-2-yl)-4-methylmorpholinium chloride DMTMM (0.75 equiv.) dissolved in 30 mL of water. The reaction was allowed to stir overnight at 56 °C, followed by filtration through cyanosilica. The filtrate was then dialyzed against DI water for over 5 days, followed by lyophilization at −80 °C. Incorporation of the small molecules was confirmed through elementary analysis.

### Cell culture & expansion

Unless otherwise specified, all cell culture supplies were purchased from VWR by Avantor. Human retinal pigment epithelial (ARPE-19) cells were purchased from American Type Culture Collection (ATCC, CRL-2302). All cells during this study were cultured using Dulbecco’s modified Eagle’s medium (DMEM/F-12) (Cytiva SH30023.01) modified by the addition of 10% fetal bovine serum (FBS) (VWR 76236-336) and 1% antibiotic-antimycotic (Fisher 15-240-062) unless otherwise specified. Cells were refreshed as needed and passaged upon reaching 80-90% confluency using 0.25% Trypsin-EDTA (Cytiva Hyclone SH30042.02).

### Cell engineering

Expression vectors and helper plasmids were developed and purchased through VectorBuilder. ARPE-19 cells were seeded into six-well plates and transfected upon reaching ∼70-80% confluency. To transfect the cells, each well was refreshed with 2 ml of Opti-MEM serum-free medium (ThermoFisher 11058021). A 2:1 ratio of expression vector to helper plasmid was used to transfect the cells, and 2.5 μg of total DNA, combined with Lipofectamine 3000 (ThermoFisher L3000015) according to protocols provided by the manufacturer, was added to each well plate. After incubation at 37 °C for 4-6 hours, the transfection medium was replaced with fresh culture medium as described previously. After transfection, cells were selected for expression of the transgene with 2.5 µg/mL puromycin until all cells, as determined via optical microscopy in a naïve control well, were killed, and then the cells were maintained using normal cell culture techniques.

### Cell line productivity measurements

To assess per-cell productivity, cells were lifted from tissue culture plastic flasks using 0.5% Trypsin + EDTA and counted using Trypan Blue and a Countess™ 3 Automated Cell Counter. After the cell count was determined, 10,000 cells were plated into 150 µL of cell culture media in replicate in 96-well plates. After allowing cells to adhere overnight, the media was aspirated, and 200 µL of new media was added. Twenty-four hours later, the media was harvested and stored at −20 °C for subsequent quantification using a human immunoglobulin G ELISA kit (ab195215) following the manufacturer’s protocol. This concentration, the cells seeded in the plate, time, and volume of media were used to calculate pg of antibody produced per cell per day.

### Monoclonal line generation

Antibody-producing clonal lines were generated from polyclonal lines of ARPE-19 cells engineered to produce antibodies as described above, after drug selection had finished. Single cells were isolated either by counting cells using trypan blue–based counting methods and plating at 0.5 cells / well in a 96-well plate or by single-cell deposition using an F.Sight single-cell dispenser (Cytena). Post-deposition of the cells into 96-well plates, the cells were maintained in DMEM/F-12 1:1 media with the addition of 20% FBS and 1% AA. Cells were allowed to grow with regular media changes for 3-5 weeks in 96-well plates. To generate distributions of per-cell secretion, standard curves with serial dilutions of specific cell numbers of the polyclonal populations were plated in 96-well plates and allowed to adhere overnight. The following day, media on the monoclonal selection plates and the standard curves was refreshed and harvested 24 hours later to be used in ELISA analysis of mAb secretion. Then, 10% alamarblue HS (ThermoFisher A50101) was added to the plates and the standard curves, and after incubation, fluorescence was measured at 560/590 excitation/emission using a Tecan infinite 200 Pro plate reader. This allowed the per-cell secretion productivity to be calculated. To visualize the distribution of clonal productivity, frequency distributions with appropriate bin sizes were plotted into histograms and then modeled using nonlinear regression analysis, fitting to either a Gaussian or a sum of two Gaussians, depending on the fit value. This was done using GraphPad Prism software (version 10.3.1, for Windows, GraphPad Software, Boston, Massachusetts, USA, www.graphpad.com). Those wells with sufficient growth and productivity were lifted and plated in 24-well plates. This process was repeated with monitoring of cell growth via optical microscopy, passaging to larger surface area well plates, and then into flasks. Conditioned media was found to improve cell survival and for some of the later monoclonal isolations was included in the monoclonal selection growth media at 20% per volume. This media was generated by culturing WT ARPE-19 cells for ∼48 hours in contact with DMEM/F12 media with 10% FBS and 1% AA. This media was then harvested and sterile filtered through a 0.2 μm filter before addition to monoclonal cell culture media.

### Alginate preparation for encapsulation

All alginates used in this study were reconstituted from lyophilized powder-form using sterile 0.8% saline. Working solutions of unmodified SLG20 alginate (Sigma, 42000001) were dissolved at 1.4% w/v, while SLG100 (Sigma, 4202101) was dissolved at a concentration of 3% w/v. Modified alginates were resuspended at 5% w/v and then blended with the 3% SLG100 at a ratio of 7:3 of modified alginate to SLG100.

### Cell encapsulation

All encapsulated cells were prepared using the same buffer reagents and methods. In preparation for encapsulation, cells were harvested from tissue culture plates, pelleted at 500 g for 5 min, and washed three times with Ca-Free Krebs buffer (4.7 mM KCl, 25 mM HEPES, 1.2 mM KH2PO4, 1.2 mM MgSO4·7H2O, 135 mM NaCl). After washing, cells were pelleted once more, supernatant was aspirated, and the pellets were resuspended in alginate at a desired concentration.

Core-Shell capsules were made by electrospraying into a crosslinking bath. Alginate-suspended cells were loaded into a Luer-lock syringe fitted to the core-extrusion port of a coaxial needle (Ramé-Hart 100-10-COAXIAL-2218) and extruded from a syringe pump (Pump 11 Pico Plus, Harvard Apparatus) at a rate of 5 mL/hr. Cell-free alginate was loaded into a second Leur-lock syringe, attached to the shell-extrusion port of the coaxial needle, and simultaneously extruded at a rate of 5.5 mL/hr. The coaxial needle was oriented vertically, with the tip of the needle approximately 2 cm above a 150 mL bath containing barium crosslinking solution (20 mM BaCl2, 250 mM d-Mannitol, 25 mM HEPES (Gibco Life Technologies) with 0.01 v/v % Tween 20). A voltage generator attached to the coaxial needle and grounded to the crosslinking bath was used to create a voltage of ∼6.25 kV. After extrusion and crosslinking of the core-shell capsules, the capsules were collected and washed three times in HEPES buffer (25 mM HEPES, 1.2 mM MgCl2·6H2O, 4.7 mM KCl, 132 mM NaCl), followed by two washes with complete cell culture medium. Capsules were then stored in cell culture medium and incubated at 37 °C until implantation. After encapsulating, capsules were imaged with bright and dark-field microscopy to verify the homogeneity of capsule size, circularity, and the core-shell morphology. mAb production from capsules was assessed by incubating capsules in cell culture media for a fixed time period, generally ∼2 hours, then harvesting supernatant samples and quantifying mAb concentration via ELISA (Abcam ab195215) following the manufacturer’s protocol. For some mAbs, antigen-specific kits were used following the manufacturer’s protocol, as described in the results: adalimumab (Abcam ab237641), ipilimumab (Abcam ab237653), pembrolizumab (Abcam ab237652).

### Lattice macrodevice design & fabrication

Macrodevice lattices were designed with Fusion360 (Autodesk) computer-aided design software (CAD) and fabricated using stereolithography additive manufacturing (Form 4B, Formlabs). Oval macrodevices 15 mm long, 4.5 mm wide, and 2 mm tall were printed with a 50 µm layer height from BioMed Clear resin (Formlabs, RS-C2-BMCL-01), a USP class IV certified material. After printing, devices underwent post-processing steps to remove cytotoxic residues and be sterilized in preparation for cell-loading and implantation (**Supplementary Fig. 14**). Lattices were first washed with a vortex washer in 90% isopropyl alcohol for 20 min, then washed for one minute in acetone. After the acetone wash, the devices were dried at room temperature for 30 minutes, washed with soapy water, and incubated in a dry oven for 24 hours at 80 °C. Lastly, devices were sterilized with an autoclave and stored at room temperature until used.

### Qualitative live/dead staining & imaging

The spatial distribution of viable cells within the hydrogels was visualized using a live/dead viability/cytotoxicity kit (Invitrogen, L3224). Capsules or macrodevices were incubated in the staining solution, prepared in PBS according to the manufacturer’s protocol, for 30 minutes at 37°C. Following incubation, they were immediately imaged using an EVOS fluorescence microscope.

### Quantitative viability by cell counting

To determine the total number of viable cells within a capsule or macrodevice, the alginate hydrogel matrix was dissolved using an enzymatic solution. Capsules or macrodevices were incubated in a solution containing 1 mg/mL alginate lyase (Millipore Sigma, A1603) and 50 mM EDTA in PBS for 30 minutes at 37°C, with gentle agitation to ensure complete dissolution. The resulting cell suspension was mixed at a 1:1 ratio with trypan blue dye and immediately counted using a Countess II automated cell counter to determine the total number of viable cells per sample.

### Murine Implantation & retrieval surgeries

All murine studies were conducted in accordance with protocols approved by Rice University’s Institutional Animal Care and Use Committee (IACUC). Depending on the study, 6-8 week old female C57BL/6J mice (Charles River Laboratories, Strain Code #027), BCD mice (B6.129S2-Ighmtm1Cgn/J, JAX Strain #002288), RAG2 mice (B6.Cg-Rag2tm1.1Cgn/J JAX Strain #008449), or NSG mice (NOD.CgPrkdcscid Il2rgtm1Wjl/SzJ JAX Strain #005557) were used, as indicated in the results and discussion sections. Prior to surgery, mice were weighed and anesthetized with 1-4% isoflurane in oxygen. Mice were kept on a water-circulating heating pad (Adroit HTP-1500) to maintain normothermia while under anesthesia. Preoperative analgesia was administered via subcutaneous injection of EthiqaXR (1.3 mg/mL), with dosage adjusted according to individual mouse body weight. All surgical instruments were sterilized by autoclaving prior to use.

#### Intraperitoneal implantation of capsules and macrodevices

The abdominal region was shaved and sterilized by scrubbing three times with betadine and isopropanol, respectively. The first incision into the subcutaneous tissue was made by elevating the skin using forceps, and a 0.5-1 cm incision was made using surgical scissors. The peritoneal wall was then elevated using forceps, and a 0.5-1cm incision was made using surgical scissors to expose the peritoneal cavity. A modified transfer pipette with a truncated tip was utilized to implant microcapsules suspended in FBS-free and Phenol-free DMEM/F12 media into the intraperitoneal cavity. Incisions were closed using absorbable sutures with an interrupted suture pattern. Postoperative monitoring included daily evaluation of wound integrity, weight, and ambulatory function for three days.

#### Subcutaneous implantation of capsules and macrodevices

The dorsal region was shaved and sterilized by scrubbing three times with betadine and isopropanol, respectively. The incision into the subcutaneous tissue was made by elevating the skin using forceps, and a 0.5 cm incision was made using surgical scissors. In the case of capsule implantation, a modified transfer pipette with a truncated tip was utilized to administer the microcapsules, which were suspended in FBS- and Phenol-free DMEM/F12 media. In the case of macrodevice implantation, macrodevices were administered with forceps. Incisions were closed using absorbable sutures with an interrupted suture pattern. Postoperative monitoring included daily evaluation of wound integrity, weight, and ambulatory function for three days.

### Subcutaneous implantation of capsules in a non-human primate

All procedures were conducted at the Tulane National Biomedical Research Center (TNBRC), an AAALAC-accredited program, under an approved Institutional Animal Care and Use Committee (IACUC) protocol P0582 and in accordance with the Guide for the Care and Use of Laboratory Animals^66,67^. A juvenile male rhesus macaque (Macaca mulatta; 3.55 years; 5.5 kg) was used.

An Ipilimumab-producing clonal cell line (ADMC P1-B1) was encapsulated in single-wall Z2A19 capsules at ∼4×10⁷ cells/mL, approximately 4 days prior to implantation. Prior to shipment, quality control included viability assessment on lysed capsules (Countess II FL) and a verification of mAb secretion by incubating ∼2 mL of capsules in 20 mL medium for 18.3 hours followed by ELISA quantification (Abcam ab195215). Capsules were shipped overnight at ambient temperature within filtered T175 flasks, and cell viability was re-checked on receipt prior to implantation. Implantation was performed under aseptic conditions in a surgical suite. The implantation site between the shoulder blades had been previously marked via tattooing. A small mid-scapular skin incision was made, a subcutaneous pocket was created by blunt dissection, and a 2 mL dose of capsules suspended in sterile FBS- and Phenol-free DMEM/F12 media was delivered subcutaneously. The incision was closed in two layers. Standard anesthesia, monitoring, and analgesia including buprenorphine were provided per institutional veterinary practice. Post-implantation blood samples were collected on days 3, 7, 14, 21, and 28 for serum ipilimumab quantification.

### Explanted capsule imaging

After explant, retrieved capsules were washed with Krebbs buffer to remove loose cell debris, transferred to a 6-well plate, and imaged with bright- and dark-field phase contrast using an Invitrogen EVOS microscope.

### Explanted capsule productivity

Following explant and phase contrast imaging, the antibody productivity of individual capsules was measured *ex vivo*. Capsules were randomly selected from each mouse and group, and single capsules were placed into individual wells of a 96-well plate containing 200 µL of complete cell culture medium and then incubated for 2 hours under standard conditions (37°C, 5% CO₂). After incubation, the media was collected, and samples were analyzed with human IgG ELISA (ABCAM, ab195215).

### Explanted capsule collagen quantification

The extent of fibrotic overgrowth on capsules explanted after 60 days and 1 year *in vivo* was quantified using a mouse Pro-Collagen I alpha 1 (COL1A1) ELISA kit (Abcam, ab210579). The capsules were then fixed in 10% formalin for 2 hours and stored in buffer at 4°C. Immediately prior to running the ELISA, 3 capsules per replicate were pooled into a single tube with PBS and were homogenized using a BeadBug microtube homogenizer and 0.5 mm glass beads (Sigma-Aldrich D1031-05). The resulting homogenate was then analyzed via ELISA according to the manufacturer’s protocol. The total mass of COL1A1 in each sample was determined from the ELISA standard curve and then divided by the number of capsules (3) to report the final value as ng of COL1A1 per capsule.

### Immunofluorescence staining and imaging

Immediately upon explant, a subset of capsules were processed for immunofluorescence analysis. Capsules were washed three times in Krebs buffer and fixed for at least 2 hours in 4% paraformaldehyde at room temperature. After fixation, samples were washed and stored in Krebs buffer at 4°C.

For staining, capsules were permeabilized with 0.5% Triton-X100 for 15 minutes at room temperature. The samples were then washed three times in PBS and blocked with a 1% BSA solution for 1 hour at room temperature. Subsequently, samples were incubated for 1 hour at room temperature in a staining solution containing 5 µg/mL anti-α-SMA-eFluor 660 [Invitrogen, 50-9760-82], 1:100 anti-CD68 CoraLite 594 fluorophore [Proteintech, CL59425747100UL], and a DAPI nuclear counterstain (NucBlue™, 2 drops/mL; [Invitrogen, R37606]), all suspended in the 1% BSA solution. Following the staining incubation, samples were washed once with a Tween-20 solution (0.1% in PBS) and twice with PBS. Capsules were then stored in Krebs buffer prior to imaging on a Nikon A1-R confocal microscope. All capsules within a comparative set were stained simultaneously with the same antibody stocks and imaged using identical acquisition settings. For final image processing and presentation, Z-stacks from the confocal microscope were converted to 2D maximum intensity projections using ImageJ (FIJI). Scale bars were added, and the brightness for all images was uniformly increased by 10% to aid visualization, with identical adjustments applied across all comparative images.

### TZM.bl HIV-1 Neutralization Assays

The TZM-bl neutralization assay was performed as previously described^68^. Briefly, mouse serum samples were tested in duplicate wells using a primary 1:50 dilution and a 3-fold titration series. Following addition of HIV-1 Env pseudovirus, plates were incubated for 1 hour at 37° C, followed by addition of TZM.bl cells (1×10^4^/well) in the presence of 11 µg/ml DEAE-Dextran (Sigma). Wells containing cells + pseudovirus (without sample) or cells alone acted as positive and negative infection controls, respectively. Following a 48-hour incubation, plates were harvested using Promega Bright-Glo luciferase reagent (Madison, WI) and luminescence detected with a Promega GloMax luminometer. Titers are reported as the reciprocal dilution of serum that inhibited 50% or 80% virus infection (ID_50_ and ID_80_ titers, respectively). Virus pseudotyped with the envelope protein of murine leukemia virus (MuLV) was used as negative control.

### Single-Cell RNA Sequencing (scRNAseq)

Mouse intra-peritoneal washes and fat pads were procured two weeks after surgical implantation of Z2A19 or SLG20 capsules containing ADC26.2 cells, or sham surgeries. Five animals were collected for each condition, with final samples including mice with sham surgery, mice implanted with Z2A19, and mice implanted with SLG20. The peritoneal lavage, fat pads, and the explanted capsules were combined and converted into single cells by using the mouse fat protocol from the Miltenyi adipose tissue dissociation kit (catalog number: 130-105-808). Afterwards, cells were washed with phosphate-buffered saline (PBS) and were subsequently stained with Alexa Fluor® 488 Anti-CD45 antibody [MRC OX-1] (Abcam catalog number: ab256254) for 1 hour. Cells were subsequently washed with PBS and resuspended in FACS buffer. Cells were then stained with NucBlue™ (Invitrogen catalog number: R37606) to assess for viability and run on an Aria III machine to collect roughly 500,000 viable cells per condition. A Chromium instrument by 10x Genomics X was used on 20,000 cells with 20,000 reads per cell, totaling 400M reads.

### Bioinformatics of scRNAseq data

The mouse (Mus musculus) reference was used from 10x Genomics. The FastQ raw data files from sequencing of the three conditions were merged (R1-R2-I1-I2) for each sample into the 10x Genomics cloud analysis platform. The mouse reference was used to map the sample files and downloaded for further analysis using Seurat in R Studio. Processing occurred as previously performed^69,70^. Briefly, data were normalized, outlier cells removed (e.g., based on RNA features or mitochondrial content), scaled, and visualized with principal component analysis. Data from each condition were integrated into a final Seurat object, allowing analysis of the three conditions: mice with sham surgery, mice implanted with Z2A19, and mice implanted with SLG20. Cell clusters were annotated based on highly expressed genes and canonical markers found in literature for mice. Cluster and gene expression analysis were performed in R Studio, like previously done^69^ and by using Seurat software packages^71^. Gene set enrichment analysis was performed using reference sets^72^.

### Secretomics data acquisition and analysis

Engineered mAb-producing cell lines and a naïve control were encapsulated in Z2A19 modified alginate as described above, and then after ∼72 hours of acclimation, groups of capsules from each cell line were washed 4x with DPBS and then incubated for ∼48 hours in DMEM/F12 1:1 media with 10% FastGro synthetic FBS and 1% Gibco Anti-Anti media. 1 mL samples were taken and frozen at −80°C. The frozen conditioned media was concentrated in a speed vac to reduce the volume to less than 50µl. The sample was then lysed in 5% SDS buffer and digested using trypsin on S-Trap micro column (Catalog: CO_2_-micro-80, ProtiFi, NY) as per manufacturer’s protocol. The peptide concentration was measured using the Pierce™ Quantitative Colorimetric Peptide Assay (Thermo Scientific 23275). The peptides were concentrated in a speed vac and dissolved in MS loading buffer (5%methanol containing 0.1% formic acid). The samples were analyzed on a Vanquish Neo UHPLC system (Thermo Fisher Scientific, San Jose, CA) coupled to Orbitrap Eclipse (Thermo Fisher Scientific, San Jose, CA) mass spectrometer. The peptide separation was carried out on a 20 cm x 75µm I.D. analytical column (Reprosil-Pur Basic C18aq, Dr. Maisch GmbH, Germany) at a 200nl/min flow rate. The MS data was acquired for 110min gradient time in a data-dependent acquisition mode. The MS1 was done in Orbitrap (120000 resolution, scan range 375-1500m/z, 50ms Injection time), followed by MS2 in Ion Trap with HCD fragmentation. The MS raw data was searched in Proteome Discoverer software (v2.1, Thermo Scientific, San Jose, CA) with Mascot algorithm^73^ (v2.4, Matrix Science) against the human NCBI refseq database (updated 2021_12_23) along with the antibody sequences. The precursor and fragment mass tolerance were set to 20ppm and 0.5Da, respectively. Maximum missed cleavages of 2 with trypsin enzyme, dynamic modification of oxidation (M), protein N-term acetylation, deamidation (N/Q) and carbamidomethyl (C) was allowed. The peptides identified from the Mascot result file were validated with 5% false discovery rate (FDR) in Percolator^74^. The gene product inference and quantification were done with a label-free iBAQ approach using the ‘gpGrouper’ algorithm^75^. The median normalized and log10-transformed iBAQ values, as well as the unique peptide spectrum matches, were used for data analysis. As an additional measure to remove trace FBS protein contaminants from analysis, in-house RNA-Seq data from ARPE-19s were used to identify proteins that ARPE-19 cells don’t express at levels > 10 TPM, such as Albumin, and those were removed from consideration. Proteomic data analysis and visualization were performed either in Python (version 3.12), utilizing custom scripts built with pandas^76^, numpy^77^, seaborn^78^, and matplotlib^79^ libraries or in GraphPad Prism (version 10.3.1).

### Histology and histological analysis

Subcutaneous implants were explanted after 1 month by excising the implant and surrounding tissue as a single unit. Samples were fixed in 10% neutral buffered formalin for 24 h and then transferred to 70% ethanol. Tissue processing, paraffin embedding, sectioning, and hematoxylin and eosin (H&E) staining were performed by the Pathology and Histology Core (HTAP) at Baylor College of Medicine. Sections (8 µm) were imaged using a Leica M165C microscope (for tile scans) and an EVOS light microscope (at 20x magnification) to provide a high-quality overview and detailed visualization of the implant site, respectively. For quantification, 4x magnification images were acquired for each replicate (n = 3, **Supplementary Fig. 13**). Fibrotic capsule thickness was then quantified from these 4x images using ImageJ (FIJI) by taking four evenly distributed measurements per section and averaging values to obtain the mean capsule thickness per implant.

### Simulated Ipilimumab I.V. Bolus

To evaluate the performance of the Z2A19 capsule platform against standard systemic administration, a 100-day I.V. bolus concentration profile was simulated for a 0.6 mg/kg dose of ipilimumab. The 0.6 mg/kg dose was selected for this simulation to yield a peak titer (C_0_) comparable to the maximum serum drug concentrations observed in our NHP study to facilitate a direct comparison of drug retention between the encapsulated cell platform and an I.V. bolus. The simulation utilized a one-compartment pharmacokinetic model with first-order elimination, defined by the equation:

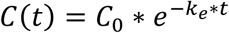

The model assumed physiological constants for a Rhesus macaque with a body weight of 5.5 kg. A weight-adjusted Volume of Distribution (V_d_) of 56 mL/kg was applied therefore resulting in a total compartment volume of 308 mL. The pharmacokinetic benchmarks used for this simulation were derived from established pre-clinical regulatory data in Cynomolgus macaques (FDA BLA 125377; EMA EPAR) due to their high degree of physiological similarity and data availability. A half-life of 8.5 days was selected to represent the non-linear clearance (target-mediated drug disposition) typically observed at low doses in NHPs. This model produced a systemic clearance of 0.19 mL/h/kg, which aligns with the prior reported range of 0.17–0.21 mL/h/kg.

### Method for calculating steady-state titer

Implant parameter ranges reflected murine-informed feasibility and implant constraints (cell density 1-8×10⁷cells/mL and per-cell productivity [PCD] 10–40 pg/cell/day).

Approximating mAb delivery through capsule and device implantation as 0^th^-order production into the blood with first-order clearance, the steady-state titer is equal to:

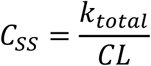

Where 𝐶_𝑆𝑆_is the steady-state concentration of mAb (ng/mL), 𝑘_𝑡𝑜𝑡𝑎𝑙_is the total production rate from the implant (ng/day), and 𝐶𝐿 is the clearance rate for the mAb being delivered (mL/day). 𝐶𝐿 is either explicitly reported in the literature or approximated using the reported half-life and estimated volume of distribution according to:

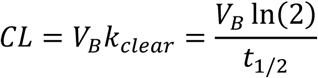

Where 𝑉_𝐵_ is the estimated volume of distribution/volume of blood (mL) and 𝑡_1/2_ is the mAb half-life (days). To relate 𝐶_𝑆𝑆_to the key implant parameters cell density (𝜎, cells/mL), cell productivity (𝑘, pg/cell/day), and volume (𝑉, mL), we substitute 𝑘_𝑡𝑜𝑡𝑎𝑙_ = 𝜎𝑘𝑉 such that:

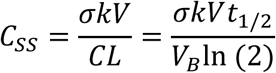

Importantly, these calculations are considered best-case scenario where cell count and clone potency remain constant *in vivo*, and all produced mAb reaches the blood.

### Statistical analysis

All statistical analyses were performed using GraphPad Prism (version 10.3.1, for Windows, GraphPad Software, Boston, Massachusetts, USA, www.graphpad.com). For >2 groups, one-way ANOVA followed by Dunnett’s T3 multiple-comparisons test (two-sided) versus the control was used; two-group comparisons used two-tailed unpaired t-tests. Reported P values are multiplicity-adjusted where applicable. Significance on graphs follows GraphPad style: ns (P>0.05), * (P≤0.05), ** (P≤0.01), *** (P≤0.001), **** (P≤0.0001).

## Acknowledgments

This work was supported by the Gates Foundation (INV-051204), and the Advanced Research Projects Agency-Health (AY1AX000003 and 1047648-490467). This material is based upon work supported by the National Science Foundation Graduate Research Fellowship Program awarded to A.D. and N.B (1842494). Any opinions, findings, and conclusions or recommendations expressed in this material are those of the authors and do not necessarily reflect the views of the National Science Foundation. HIV pseudovirus neutralization assays were performed by the lab of Michael Seaman at the Center for Virology and Vaccine Research supported by Bill and Melinda Gates Foundation grant INV-036842. Mass spectrometry and analysis services were provided by the BCM Mass Spectrometry Proteomics Core, with contributions by Antrix Jain, Alexander Saltzman, and Anna Malovannaya, with support by (RRID:SCR_027015) the Dan L. Duncan Comprehensive Cancer Center Award (P30 CA125123), and CPRIT Core Facility Awards (RP210227). Histology services were provided by the Pathology and Histology Core (HTAP) at Baylor College of Medicine, which is supported by the P30 Cancer Center Support Grant (NCI-CA125123). This project was supported in part by the Cytometry and Cell Sorting Core at Baylor College of Medicine with funding from the CPRIT Core Facility Support Award (CPRIT-RP240432) and the NIH (CA125123, OD036336, and OD038251). This project was supported in part by the Genomic and RNA Profiling Core at Baylor College of Medicine. The NHP study was performed at the Tulane National Biomedical Research Center (RRID: SCR_008167). We thank the Division of Veterinary Medicine at TNBRC, particularly Dr. Katherine A. Johnson, for surgical expertise and veterinary care during the NHP study. Confocal microscopy and additive manufacturing resources were supported in part by the Shared Equipment Authority at Rice University. Schematics & renders were created using Biorender.com.

## Author Contributions

C.F., A.D., S.P., M.D., O.V. designed the studies and interpreted the results. A.D. designed genetic constructs and developed engineered cell lines. S.P., C.S., P.B. performed the synthesis. C.F. designed and developed retrievable macrodevices. C.F., A.D., S.P., Z.W., N.B., D.M., I.M. conducted mouse studies. E.H., S.C., A.A. conducted the NHP study. C.F., A.D., Z.W., Y.K. performed *in vitro* assays. C.F., A.D., S.P., J.D. analyzed and visualized data. T.G. performed scRNA-Seq bioinformatics analysis. C.L., M.S. performed TZM.bl neutralization assays. C.F., A.D., S.P. wrote the manuscript. C.F., A.D., S.P., T.G., A.A., O.I., R.G., M.D., O.V. provided editing and revision feedback. All authors reviewed and approved the final manuscript.

## Extended Data

**Extended Data Figure 1.**
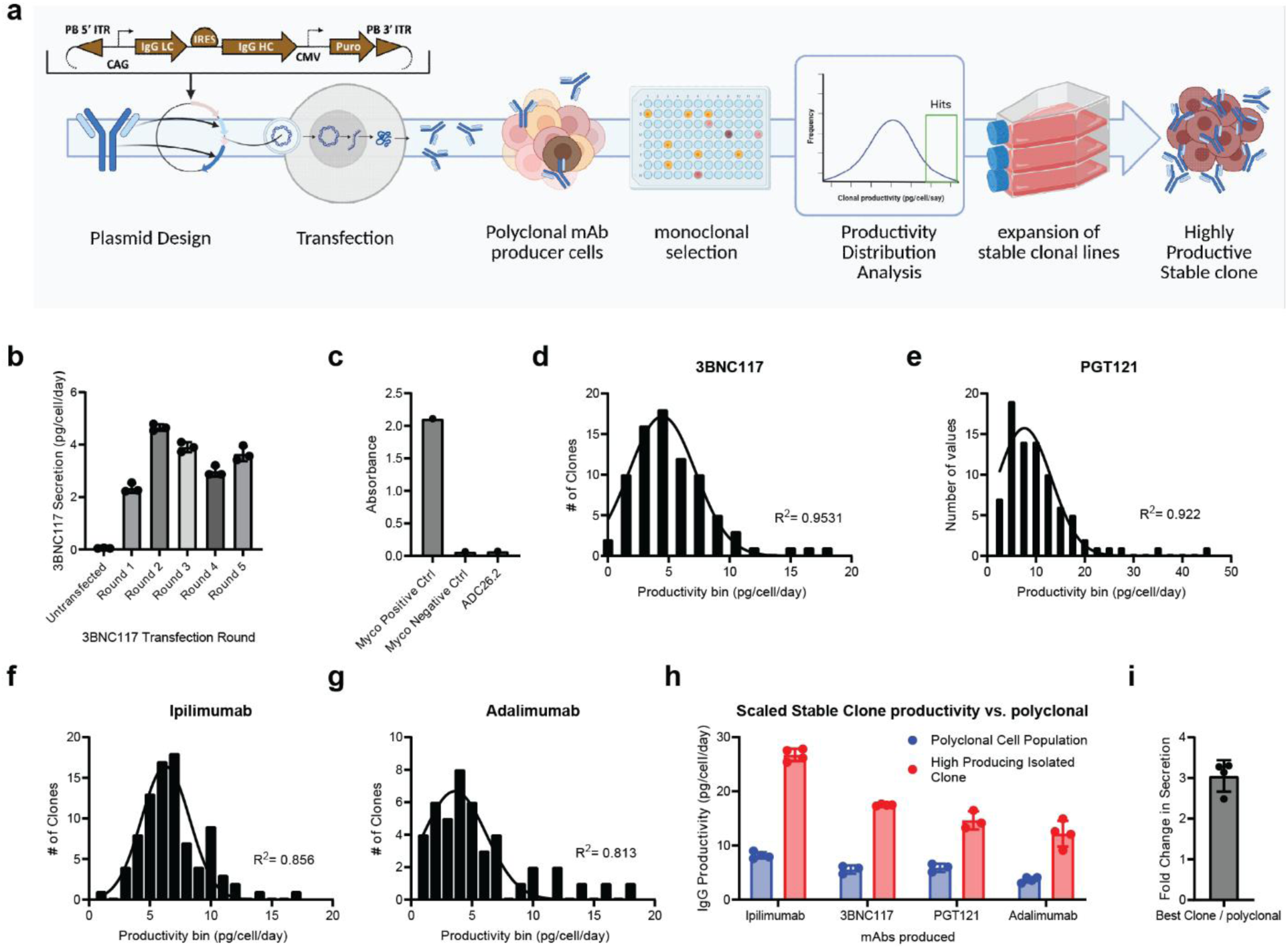
Extended cell engineering and monoclonal selection pipeline. **a,** Schematic illustrating pipeline **b,** Per cell secretion across 5 rounds of transfection into ARPE-19 cells with 3BNC117 producing plasmid and an un-transfected control. **c,** One example of mycoplasma testing performed routinely on cultured cell lines **d-g,** per cell secretion frequency distributions of clonal populations screened while in 96-well plates, with Gaussian fits shown (panel d data also shown in Fig 1c main text). **h,** Highest secreting scaled stable isolated clonal populations compared to pre-monoclonal selection polyclonal population. **i,** Data showing fold change in secretion of selected monoclonal populations compared to their respective polyclonal populations.

**Extended Data Figure 2.**
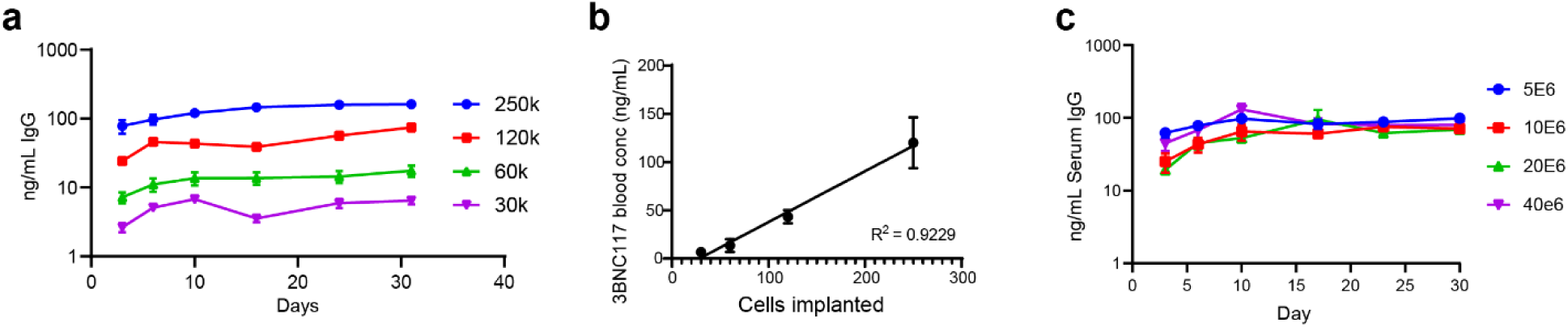
Control of monoclonal antibody dosing from implanted capsules. **a,** Serum mAb levels over time with increasing implanted cell doses in NSG mice. **b,** Linear correlation between serum concentration and implanted cell dose at Day 10 from panel A. **c,** Time course of serum titers in NSG mice over 30 days with equalized cell doses across groups in which capsules were made with different cell densities. Implanted capsule number was adjusted to achieve the same effective dose.

**Extended Data Figure 3.**
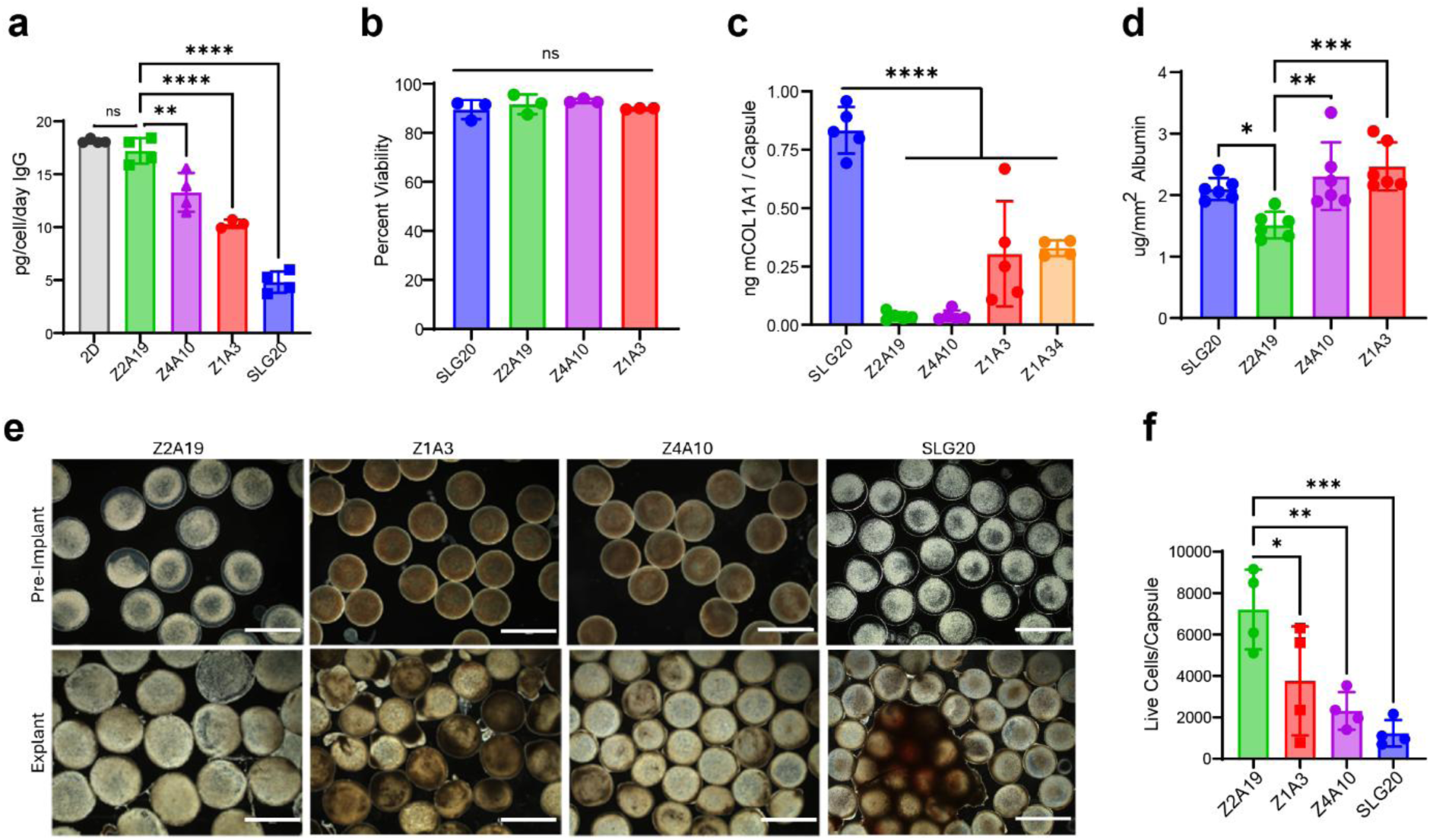
Extended assessment of modified alginate capsules. **a,** Per day productivity of encapsulated cells compared to unencapsulated cells, n=4 for all groups other than Z1A3 (n=3). **b,** Percent viability of cells encapsulated in modified and unmodified alginate (n=3). **c,** Collagen overgrowth (fibrosis) quantified at explant (±SD, n=4 for Z1A34, n=5 for all other groups). **d,** In vitro albumin adsorption on modified and unmodified capsules (SD, n=5). **e,** 1-year explanted capsule images. Scale bars = 2 mm. **f,** Live cells per explanted capsule from n=4 mice per condition.

**Extended Data Figure 4.**
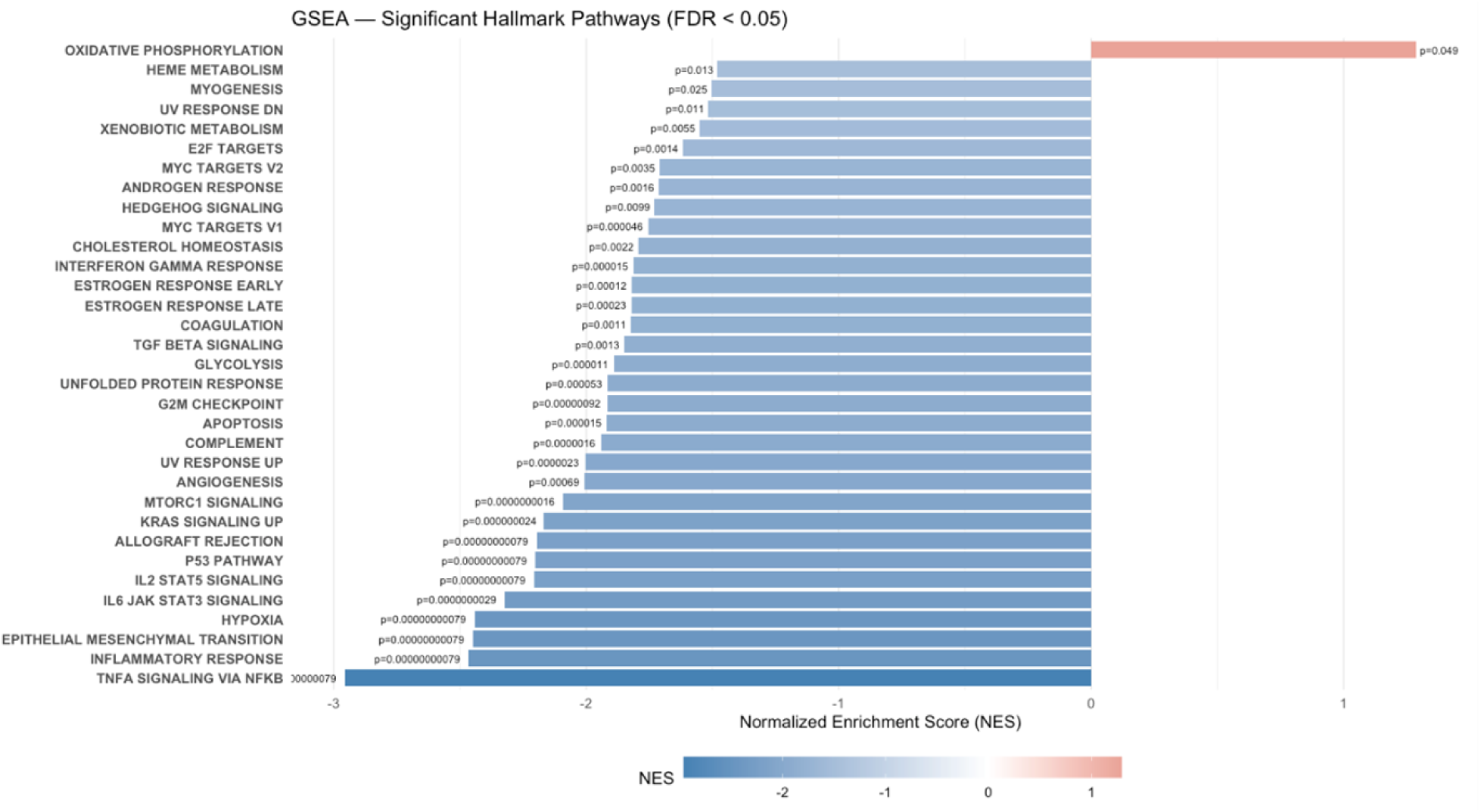
Significant Hallmark Pathways. Bar chart with annotated P-Values showing the Normalized Enrichment Score from scRNA-Seq data of Z2A19 relative to SLG20, only those with FDR < 0.05 are shown.

**Extended Data Figure 5.**
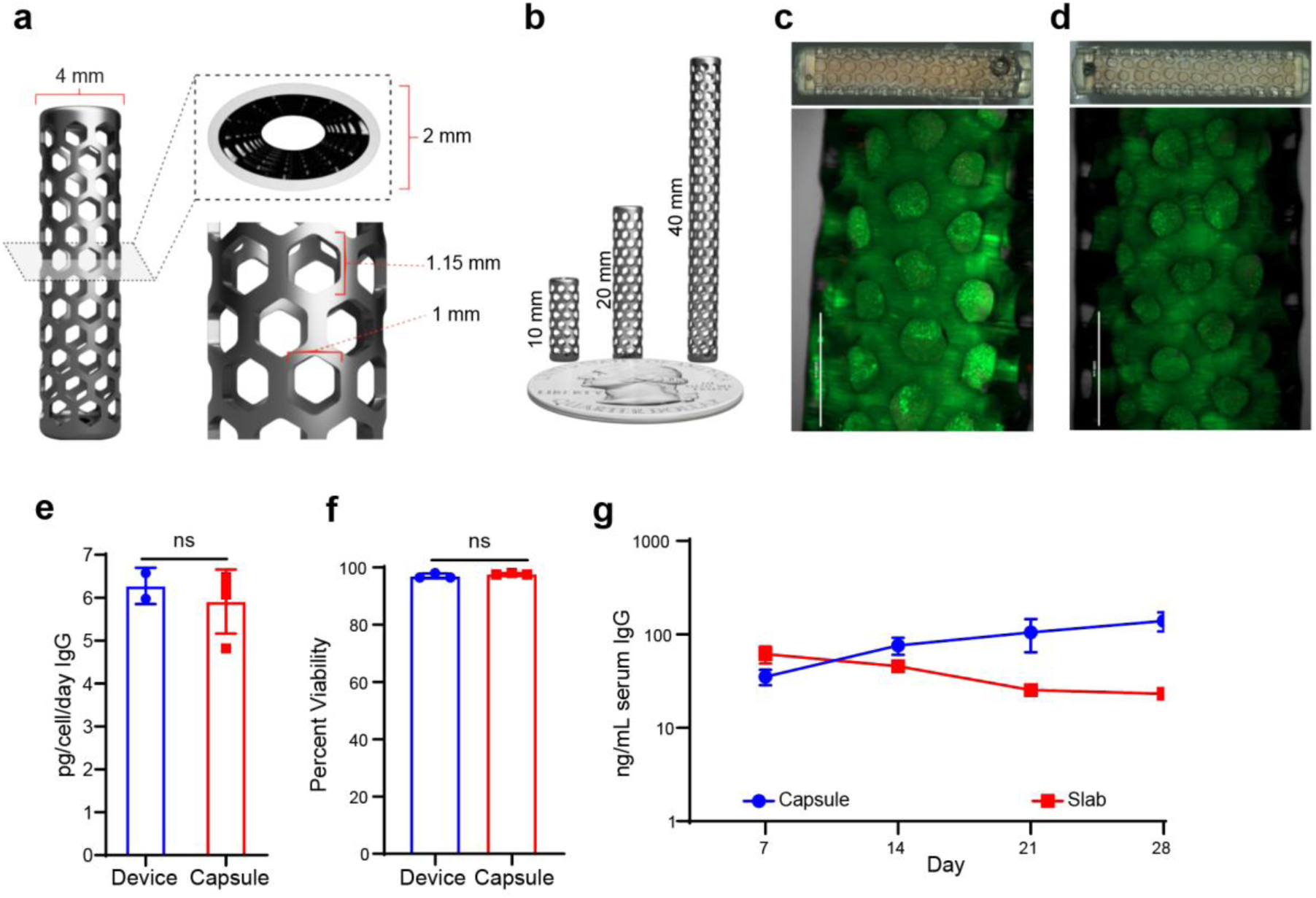
Device design and development. **a,** Lattice profile and hexagonal pore dimensions. **b,** Device length can be increased to scale loading volume without changing core lattice and overall profile features. **c-d,** Live-dead images and bright field images of devices loaded with either a **c**, solid alginate slab or **d**, alginate capsules. **d-e,** In-vitro, slab and capsule loaded devices demonstrate identical productivity and viability. **f,** In-vivo, capsule-loaded devices outperform devices containing a slab of alginate.

**Extended Data Figure 6.**
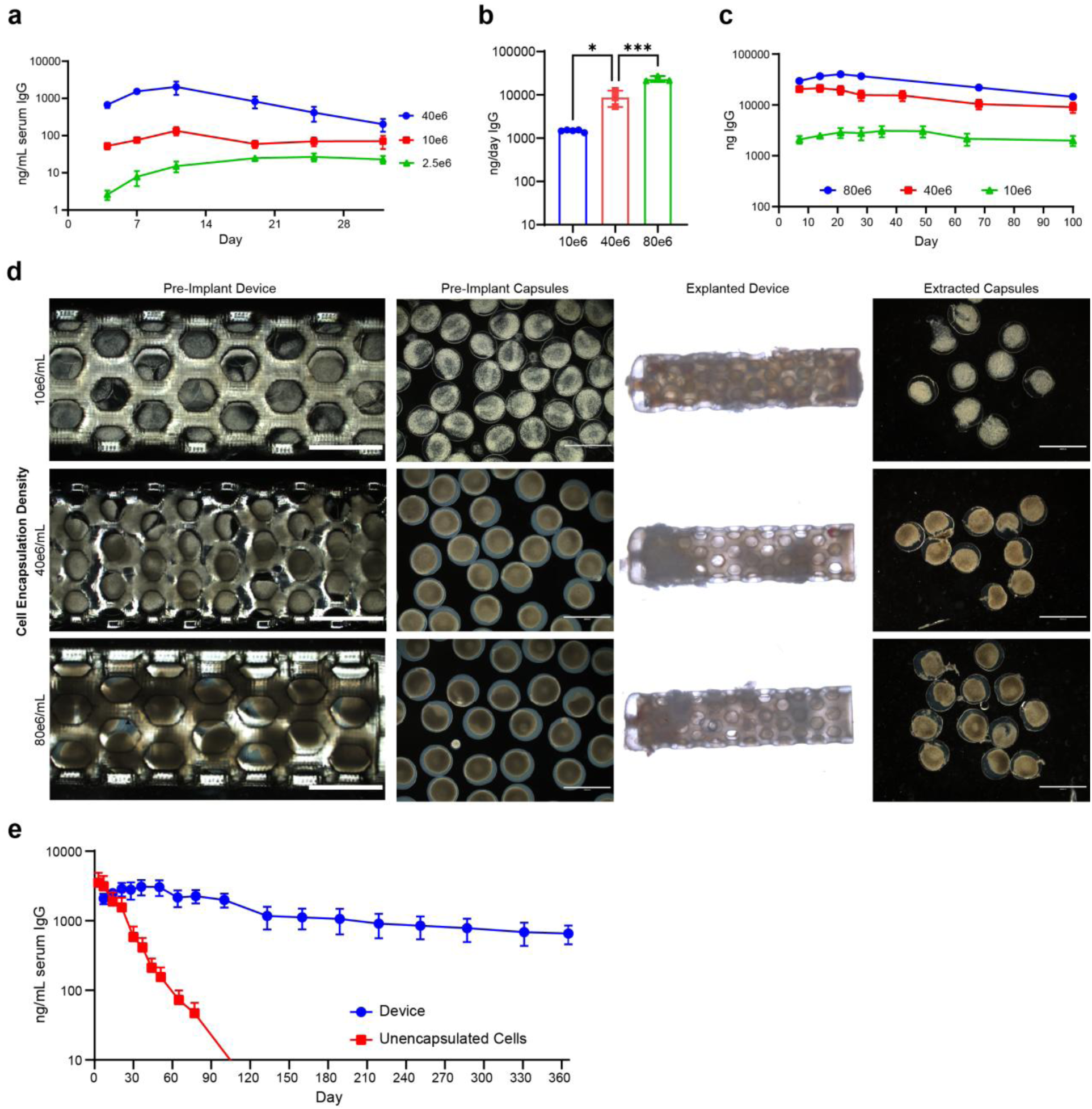
Extended in vivo device data. **a,** Dose titration of devices containing Z4A10 capsules implanted in the IP compartment of NSG mice. Device size remained constant while encapsulation density varied between 2.5-40e6 cells/mL to adjust the number of cells within each device. **b,** Pre-implant productivity of Z2A19-capsule loaded devices prior to implanting in BCD mice (panel c). Device size remained constant while encapsulation density varied between 10-80e6 cells/mL to change the number of cells within each device. **c,** Dose titration of devices containing Z2A19 capsules implanted in the IP compartment of BCD mice. **d,** Images of devices and Z2A19 capsules contained within the devices before implant and after explant after 100 days in BCD mice. Capsules extracted from explanted devices are strikingly free of cellular overgrowth. **e,** 1-year stable titer from Z2A19 capsule loaded devices implanted in the IP compartment of BCD mice compared to unencapsulated cells injected into the same compartment. Titer data in panel e is extended data from the Figure 4 panel d IP implant group.

**Extended Data Figure 7.**
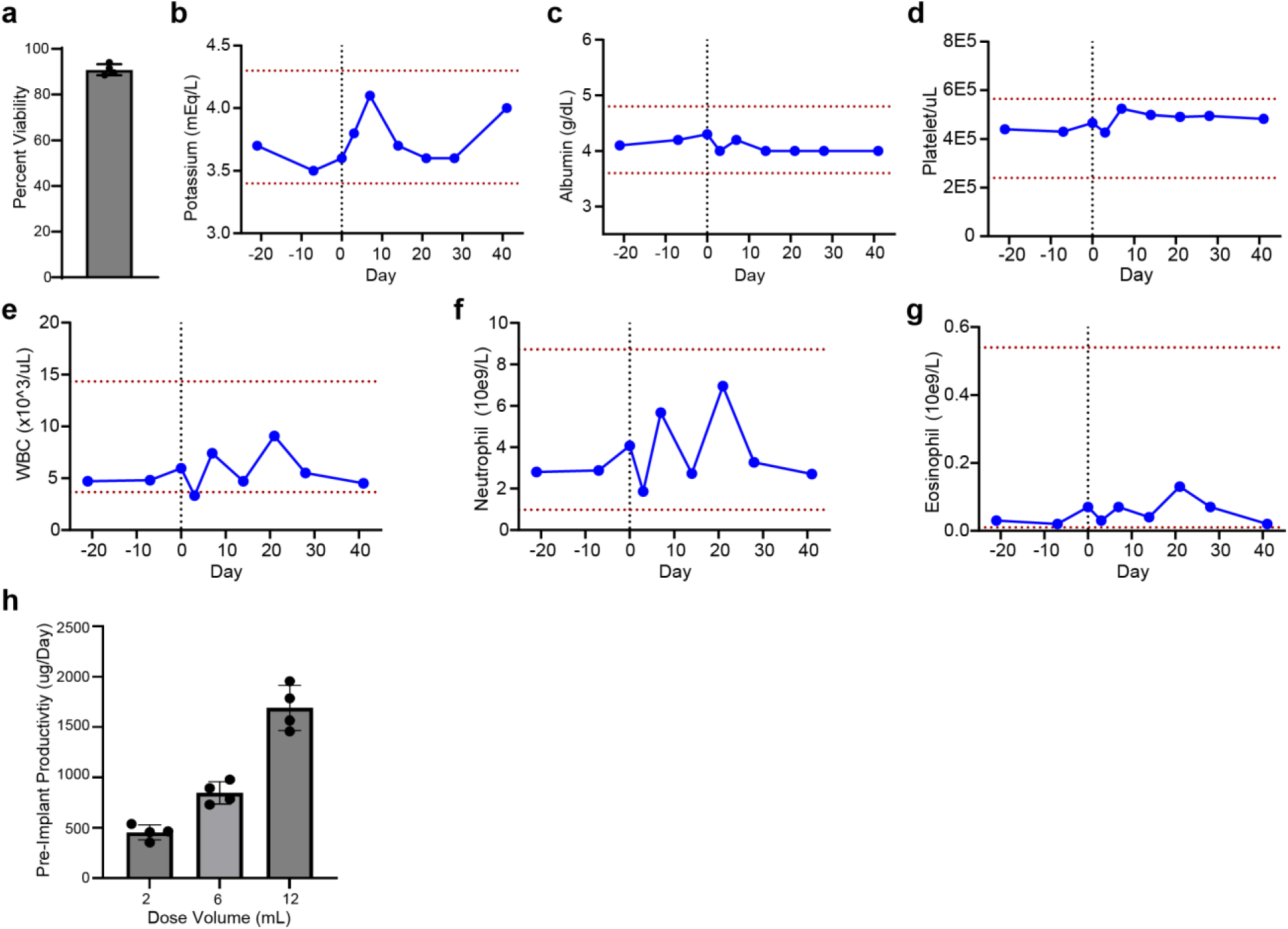
Extended non-human primate study data. **a,** Percent viability of encapsulated cells prior to implant (90.8%). **b-g,** NHP blood chemistry and CBC levels. **b,** Potassium levels. **c,** Albumin levels. **d,** Platelet counts. **e,** White blood cell counts. **f,** Neutrophil counts. **g,** Eosinophil counts. **h,** Pre-implant NHP dose productivity.

