## Supplementary Information for "Once yearly cell-based therapy for sustained and dose tunable delivery of monoclonal antibodies"

#### Supplementary Tables

***Table 1.*** *Source and catalog numbers of materials, tools, and equipment used throughout this manuscript.*

| Reagent/Resource | Source/Company | Catalog Number | Additional Information |
|---|---|---|---|
| ARPE-19 cells | ATCC | CRL-2302 | Human retinal pigment epithelial cells |
| DMEM/F-12 medium | Cytiva | SH30023.01 | 1:1 formulation |
| Fetal bovine serum (FBS) | VWR | 76236-336 | 10% in culture medium |
| Antibiotic-antimycotic | Fisher | 15-240-062 | 1% in culture medium |
| Trypsin-EDTA (0.25%) | Cytiva Hyclone | SH30042.02 | For passaging |
| Opti-MEM medium | ThermoFisher | 11058021 | Serum-free |
| Lipofectamine 3000 | ThermoFisher | L3000015 | Transfection reagent |
| Puromycin | Gibco | A1113803 | 2.5 μg/mL for selection |

| | | | |
|---|---|---|---|
| alamarBlue HS | ThermoFisher | A50101 | 10% solution for viability |
| SLG20 alginate | Sigma | 42000001 | Unmodified alginate |
| SLG100 alginate | Sigma | 4202101 | Unmodified alginate |
| Coaxial needle | Ramé-Hart | 100-10-COAXIAL-2218 | For core–shell capsule fabrication |
| Syringe pump | Harvard Apparatus | Pump 11 Pico Plus | For electrospraying capsules |
| $BaCl_2$ crosslinking bath chemicals | Gibco Life Technologies | — | 20 mM $BaCl_2$, 250 mM d-Mannitol, 25 mM HEPES, 0.01% Tween-20 |
| Human IgG ELISA kit | Abcam | ab195215 | For quantifying antibody |
| Adalimumab ELISA kit | Abcam | ab237641 | Antigen-specific |
| Ipilimumab ELISA kit | Abcam | ab237653 | Antigen-specific |
| Pembrolizumab ELISA kit | Abcam | ab237652 | Antigen-specific |
| COL1A1 ELISA kit | Abcam | ab210579 | Pro-Collagen I alpha 1 assay |
| MycoProbe Mycoplasma Detection Kit | R&D Systems | CUL001B | For regular myco-plasma testing of cells in culture |
| 3D printer (Form 4B) | Formlabs | — | Stereolithography printer |
| BioMed Clear resin | Formlabs | RS-C2-BMCL-01 | For 3D-printed macrodevice lattice |
| Live/Dead Viability Kit | Invitrogen | L3224 | For cell viability staining |
| Alginate lyase | Millipore Sigma | A1603 | For hydrogel dissolution |
| Countess III Automated Cell Counter | ThermoFisher | AMQAX2000 | For viability counts |
| C57BL/6J mice | Charles River | 027 | Immunocompetent strain |
| BCD mice | JAX | 002288 | B6.129S2-Ighmtm1Cgn/J |
| RAG2 mice | JAX | 008449 | B6.Cg-Rag2tm1.1Cgn/J |
| NSG mice | JAX | 005557 | NOD.Cg-Prkdcscid Il2rgtm1Wjl/SzJ |
| EthiqaXR | Fidelis | — | Buprenorphine extended-release analgesic, 1.3 mg/mL |
| Heating pad | Adroit | HTP-1500 | For surgical warming |
| Anti-α-SMA-eFluor 660 | Invitrogen | 50-9760-82 | For immunofluorescence |
| Anti-CD68 CoraLite 594 | Proteintech | CL59425747100UL | For immunofluorescence |
| NucBlue (DAPI) | Invitrogen | R37606 | Nuclear counterstain |
| Anti-CD45 Alexa Fluor 488 | Abcam | ab256254 | Flow cytometry marker |
| NucBlue (DAPI) | Invitrogen | R37606 | Nuclear counterstain |
| Miltenyi adipose dissociation kit | Miltenyi | 130-105-808 | For single-cell prep |
| S-Trap micro column | ProtiFi | CO2-micro-80 | For proteomics digestion |
| Pierce peptide assay | Thermo Scientific | 23275 | For proteomics peptide quantification |
| Orbitrap Eclipse MS | Thermo Scientific | — | Mass spectrometer |
| Vanquish Neo UHPLC system | Thermo Fisher | — | Proteomics LC system |
| 10x Genomics Chromium | 10x Genomics | — | Single-cell capture |
| Seurat | Open source | — | Bioinformatics software (R package) |
| EVOS fluorescence microscope | Invitrogen | — | For cell imaging |
| Nikon A1-R confocal microscope | Nikon | — | For immunofluorescence |
| F.Sight single-cell dispenser | Cytena | — | For monoclonal isolation |

***Table 2.*** *Top 20 proteins across secretome sample groups and their function not including mAbs*

| Gene ID | UniProt ID | UniProt Function Summary |
|---|---|---|
| CTSD | P07339 | Acid protease active in intracellular protein breakdown. Plays a role in APP processing following cleavage and activation by ADAM30 which leads to APP degradation |
| CST3 | P01034 | Cysteine protease inhibitor; regulates cathepsin activity |
| GSN | P06396 | Actin filament severing/capping protein; regulates cytoskeleton dynamics |
| SERPINF1 | P36955 | Neurotrophic protein; induces extensive neuronal differentiation in retinoblastoma cells. Potent inhibitor of angiogenesis.it exhibits no serine protease inhibitory activity. |
| RARRES2 | Q99960 | A component of desmosome cell-cell junctions which are required for positive regulation of cellular adhesion |
| B2M | P61769 | Component of the class I major histocompatibility complex (MHC). Involved in the presentation of peptide antigens to the immune system. |
| SPARC | P09486 | Appears to regulate cell growth through interactions with the extracellular matrix and cytokines. Binds calcium, copper, collagen, albumin, thrombospondin, PDGF and cell membranes |
| EFEMP1 | Q12805 | Binds EGFR, the EGF receptor, inducing EGFR autophosphorylation and the activation of downstream signaling pathways. May play a role in cell adhesion and migration. |
| PPIA | P62937 | Catalyzes the cis-trans isomerization of proline imidic peptide bonds in oligopeptides. Activates endothelial cells (ECs) in a pro-inflammatory manner |
| IGFBP7 | Q16270 | Binds IGF1 and IGF2 with a relatively low affinity. Stimulates prostacyclin (PGI2) production. Stimulates cell adhesion. Acts as a ligand for CD93 to play a role in angiogenesis |
| LGALS1 | P17931 | β-galactoside-binding lectin; mediates cell–cell and cell–matrix adhesion |
| ECM1 | Q16610 | ECM structural/regulatory protein; Stimulates the proliferation of endothelial cells and promotes angiogenesis. Inhibits MMP9 proteolytic activity. |
| NPC2 | P61916 | Lysosomal cholesterol transporter; essential for lipid metabolism |
| TIMP1 | P01033 | Inhibits matrix metalloproteinases; regulates ECM turnover |
| CLU | P10909 | Molecular chaperone; involved in apoptosis, lipid transport, and complement regulation |
| PSAP | P07602 | Precursor of saposins; essential for sphingolipid degradation in lysosomes |
| IGFBP6 | P24592 | Binds IGFs, preferentially IGF-II; inhibits growth, Activates the MAPK signaling pathway and induces cell migration |
| PTGDS | P41222 | Prostaglandin D synthase; produces $PGD_2$ (modulates sleep, inflammation) |
| DKK3 | Q9UBP4 | Wnt signaling inhibitor |
| TGFBI | Q15582 | ECM adhesion molecule induced by TGF-β; influences cell adhesion |
| FN1 | P02751 | Fibronectin; cell adhesion, migration, and wound healing |
| CHI3L1 | P36222 | Chitinase-3-like protein 1; tissue remodeling, inflammation |
| C3 | P01024 | Complement component 3; activates complement system |

| | | |
|---|---|---|
| TIMP2 | P16035 | Metalloproteinase inhibitor; ECM homeostasis |
| CALR | P27797 | Calreticulin; Calcium-binding chaperone that promotes folding, oligomeric assembly and quality control in the endoplasmic reticulum (ER) via the calreticulin/calnexin cycle. |
| CP | P00450 | Ceruloplasmin; ferroxidase activity, iron homeostasis |
| SOD3 | P08294 | Extracellular superoxide dismutase; ROS regulation |
| MMP2 | P08253 | Matrix metalloproteinase-2; ECM remodeling, angiogenesis |

***Table 3.*** *Reported Clearance Rates and Half-Lives for mAbs of Interest*

| mAb | CL | Source |
|---|---|---|
| 3BNC117 | 390 mL/day | Approximated based on reported half-life in HIV+ patients (9 days) by Caskey et al. [1] and a blood volume of 5000 mL. |
| PGT121 | 250 mL/day | Approximated based on reported-half-life for HIV+ patients on ART (14 days) reported by Stephenson et al. [2] and a blood volume of 5000 mL. |
| Ipilimumab | 400 mL/day | Ipilimumab FDA label |
| Pembrolizumab | 200 mL/day | Pembrolizumab FDA label |
| Rituximab | 340 mL/day | Rituximab FDA label |
| Adalimumab | 290 mL/day | Adalimumab FDA label. |

### Supplementary Results

#### Supplementary cell engineering figures

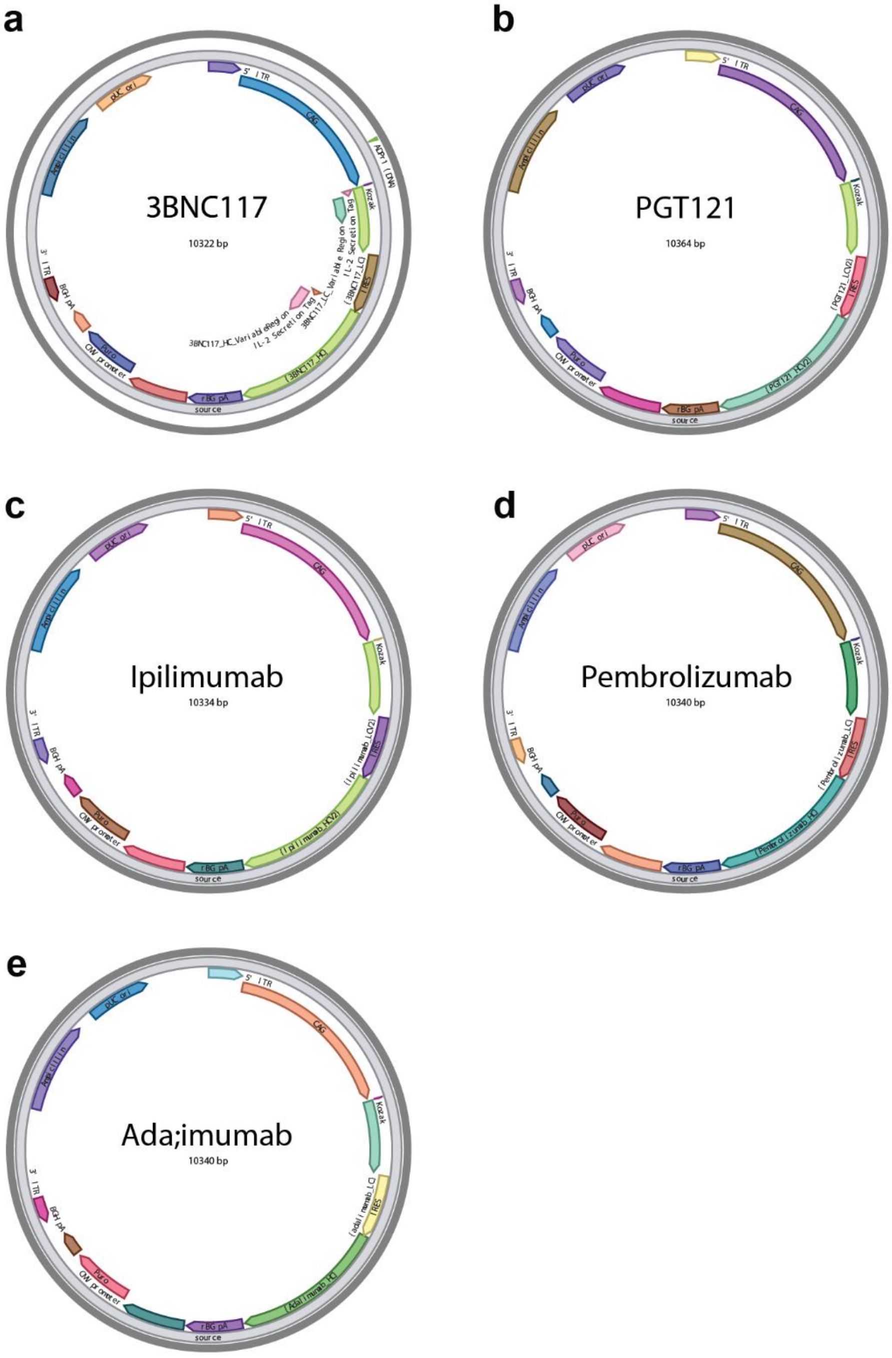

**Supplementary Figure 1. Plasmid maps for mAb expression**. **a-e**, Genetic constructs used to engineer cells to produce mAbs of interest 3BNC117, PGT121, Ipilimumab, Pembrolizumab, Adalimumab respectively.

### Supplementary capsule optimization and in vivo figures

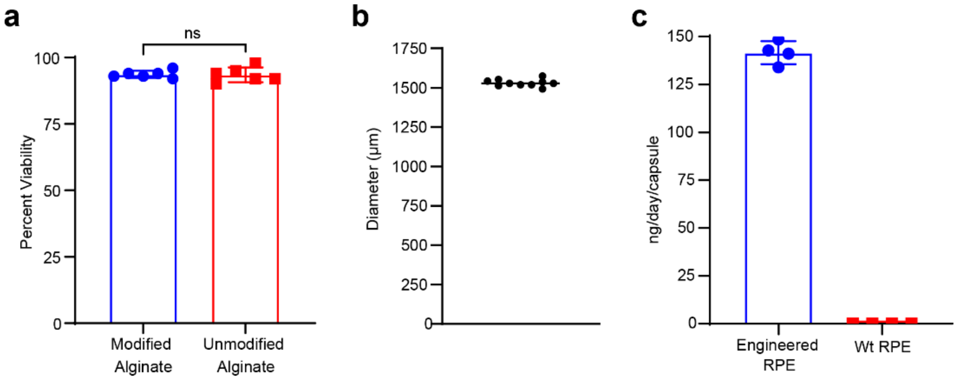

**Supplementary Figure 2: Encapsulated cell quality control. a,** Cell viability between modified and unmodified Alginate as determined by trypan blue staining (n=6). **b,** Capsule diameter distribution (n=10). **c,** IgG secretion rate from capsules engineered cells vs naïve ARPE-19s (n=4).

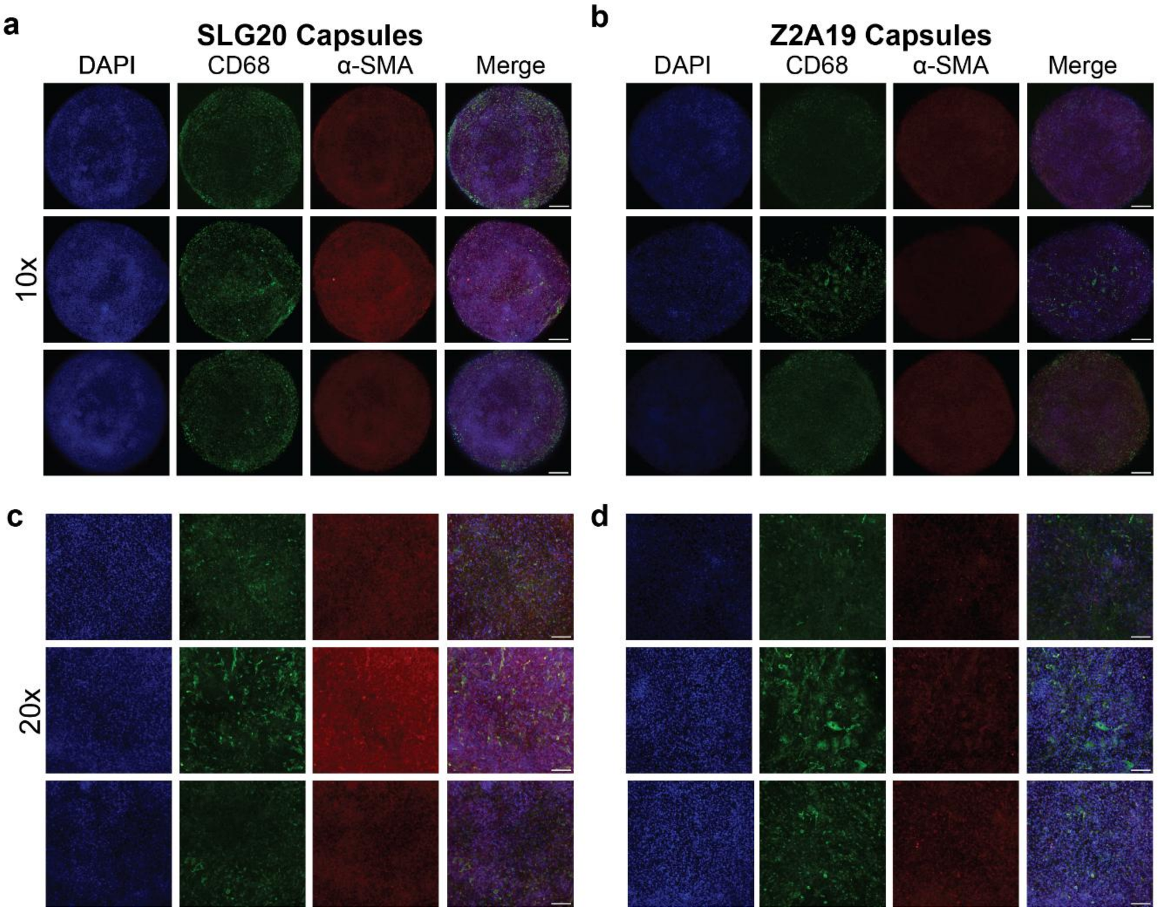

**Supplementary Figure 3. Immunofluorescence images of unmodified SLG20 and Z2A19 modified capsules**. Capsules were explanted from C57BL/6 mice (n=3) for imaging after 2-weeks and stained with DAPI nuclear stain, anti-CD68 to visualize macrophages, and anti-a-SMA to visualize myofibroblasts. **a-b**, show macroscale images of stitched and z-stacked capsules using a 10x objective. **c-d**, show z-stacked images of capsules obtained with a 20x objective. Scale bar for 10x images = 200 μm. Scale bar for 20x images = 100 μm.

#### scRNA-Seq supplementary figures

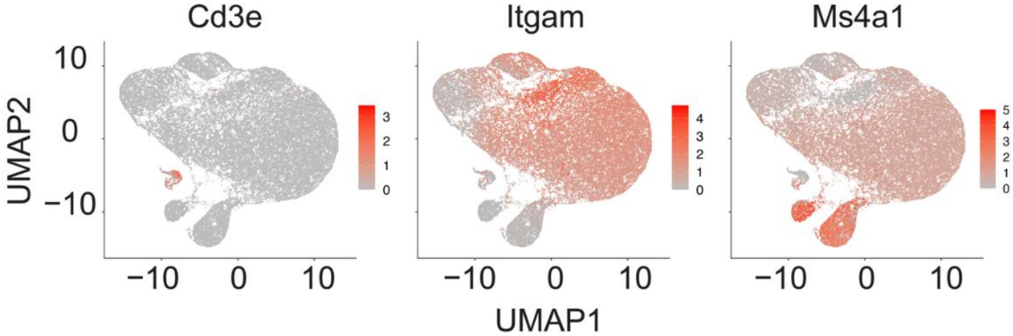

**Supplementary Figure 4. UMAP of cluster-specific genes for myeloid cells, B cells, and T cells**. Integrated clusters across three conditions: sham mice, mice implanted with Z2A19 material with cells, and mice implanted with SLG20 material with cells. Genes Cd3e, Itgam, and Ms4a1 are plotted for specific types of cell types (T cells, myeloid cells, and B cells respectively). Red dots indicate cells with higher gene expression while gray dots represent cells with lower gene expression.

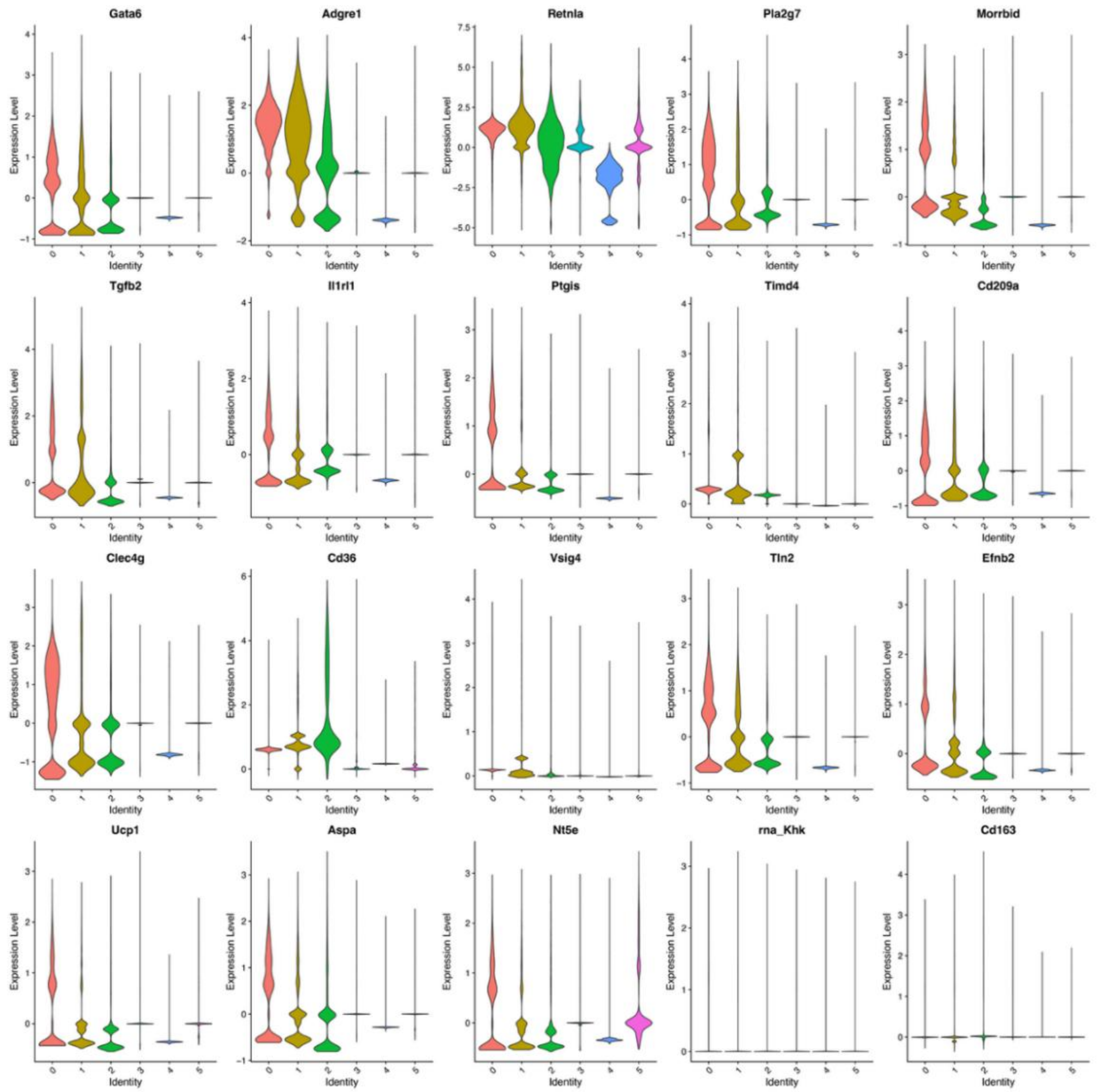

**Supplementary Figure 5. Anti-inflammatory macrophage markers.** Violin plot of genes associated with anti-inflammatory macrophages. Numbers on the x-axis are the cluster number.

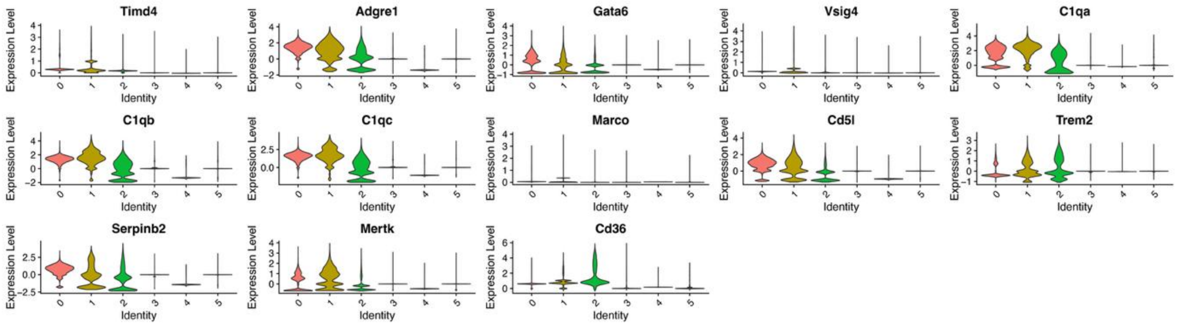

**Supplementary Figure 6. Supplementary Figure 6: Resident macrophage markers**. Violin plot of genes associated with resident macrophages. Numbers on the x-axis are the cluster number.

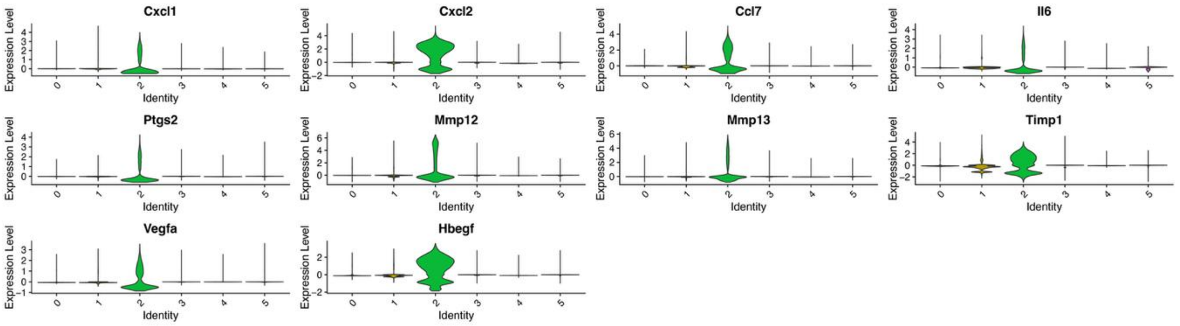

**Supplementary Figure 7: Inflammatory macrophage markers.** Violin plot of genes associated with inflammatory macrophages. Numbers on the x-axis are the cluster number.

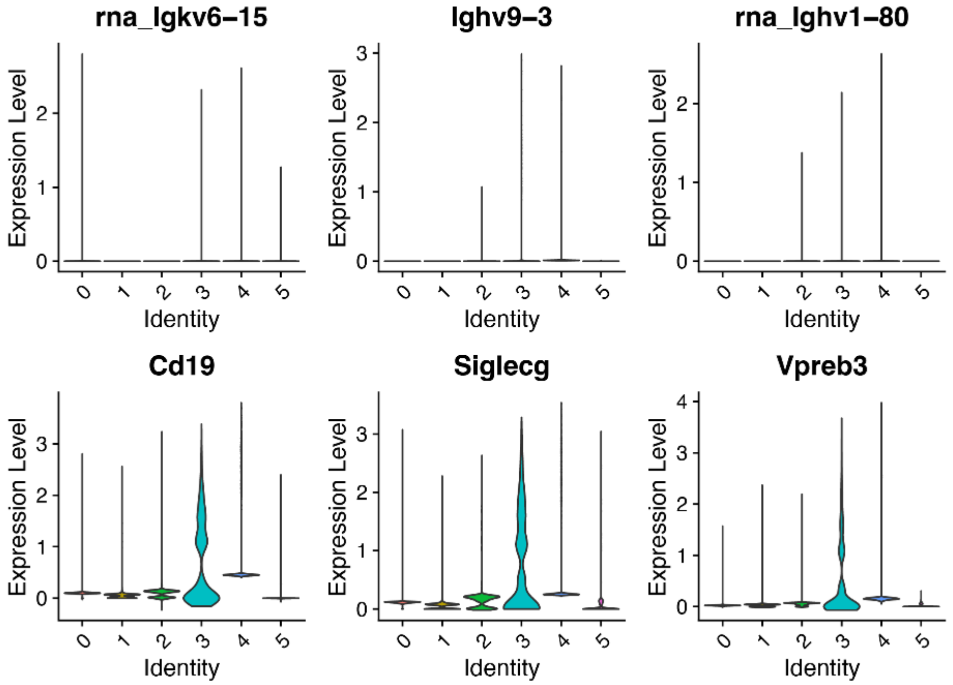

**Supplementary Figure 8: Follicular B Cell markers** Violin plot of genes associated with Follicular B cells. Numbers on the x-axis are the cluster number.

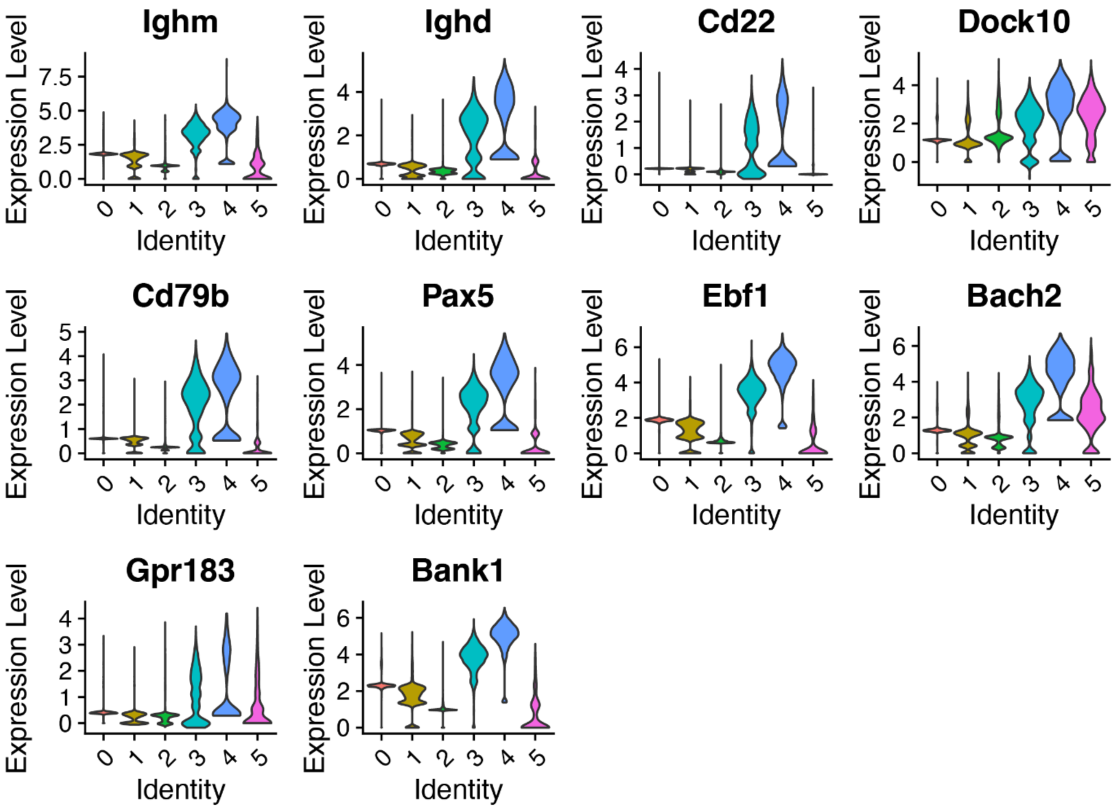

**Supplementary Figure 9: Memory B Cell markers.** Violin plot of genes associated with Memory B cells. Numbers on the x-axis are the cluster number.

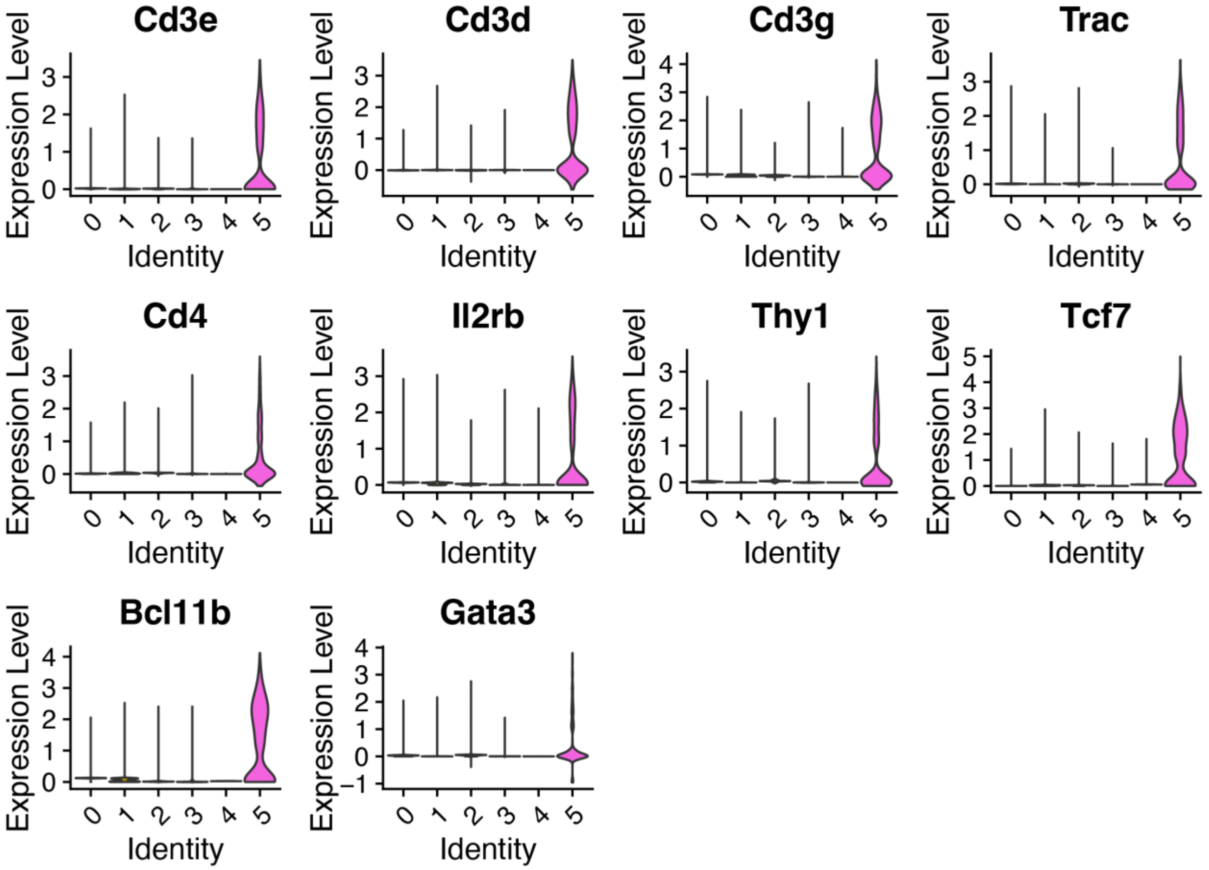

**Supplementary Figure 10. CD4 T Cell markers.** Violin plot of genes associated with CD4 T cells. Numbers on the x-axis are the cluster number.

#### Proteomics & additional mAb supplementary figures

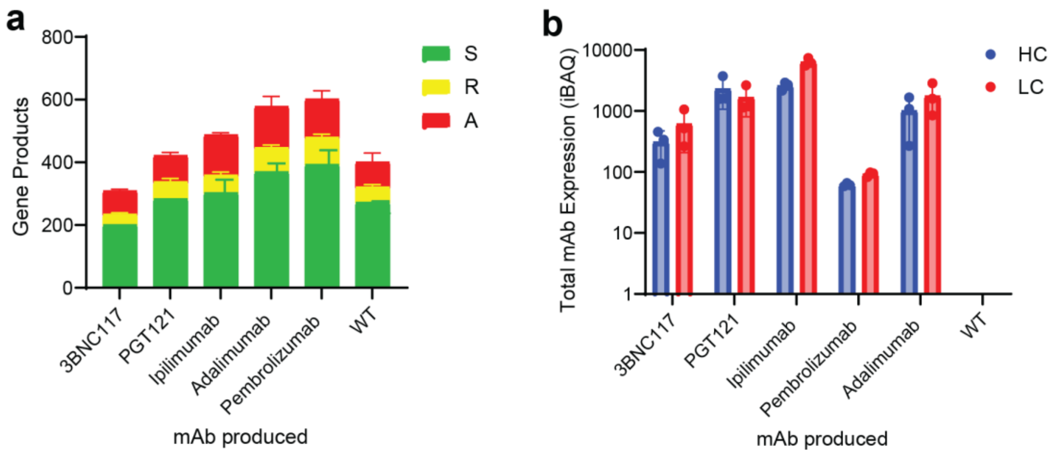

**Supplementary Figure 11: Secretomics additional data. a,** Identified protein quality across identified gene products **b,** iBAQ scores across mAbs across sample groups.

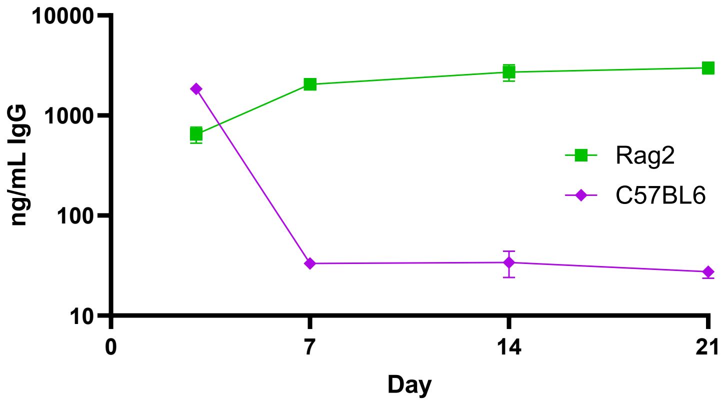

**Supplementary Figure 12. Differences in mAb titer and pharmacokinetics in immunocompetent vs immunocompromised mice.** Time vs Serum levels of Adalimumab secreting Implants in C57BL6 and RAG2 mice (n=5).

#### Supplementary H&E Images

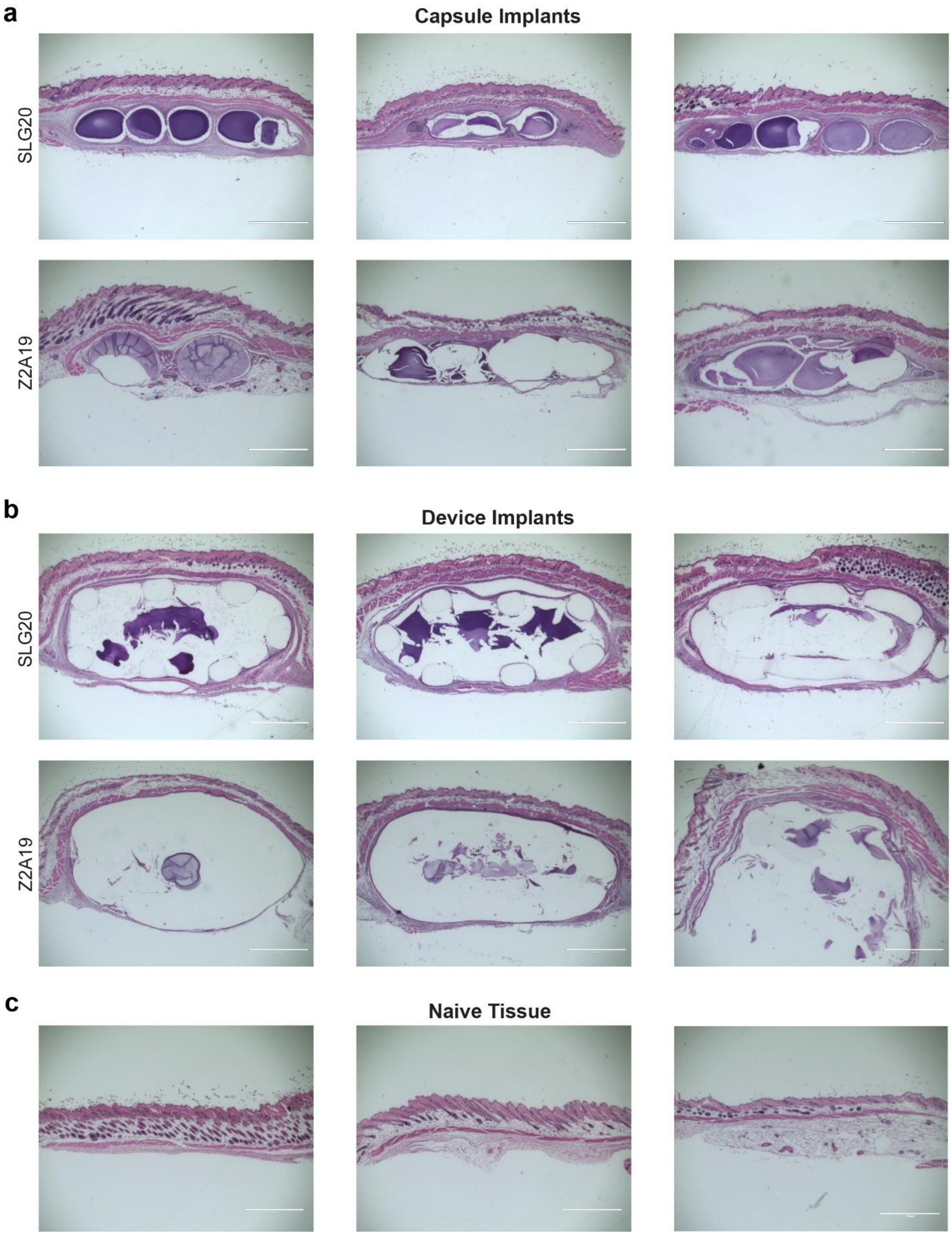

**Supplementary Figure 13. Extended H&E images.** 4x magnification Images of H&E-stained subcutaneous tissue harvested from C57BL/6 mice implanted with SLG20 or Z2A19 capsules (**A**) or macrodevices (**B**) for 1 month (n=3). Images of Naïve subcutaneous tissue (**C**) are shared for reference. These images were used for fibrotic capsule thickness quantification in figure 4 panels J and K. Scale bars = 1mm.

### Supplementary Methods

#### Small molecule synthesis

All the small molecules were synthesized using click chemistry. The following synthesis route was followed for the synthesis of all the molecules except B2A17. Briefly, to a two necked round bottom flask containing 180 mL of water and methanol in 5:1 ratio under nitrogen was added alkyne followed by addition of tris((1-benzyl-4- triazolyl)methyl) amine (TBTA) (0.25 equiv.), followed by addition of triethylamine (0.25 eq) and copper(I) iodide (0.1 eq) The reaction was then allowed to cool down to 0°C for 15 minutes while still purging the nitrogen. Respective peg- linkers (Azido-PEG-amine) were then added at 1.1 eq to the reaction mixture, and it was allowed to stir for another 5 mins at room temperature followed by heating at 55°C overnight. The reaction was then filtered over celite, solution was evaporated using rotovap and the crude was purified using liquid chromatography with dichloromethane: ultra (22% MeOH in DCM with 3% NH4OH) 0% to 40% on a 120 gm ISCO silica column.

B2A17 synthesis: 3-iodobenzylamine (1 equiv.) was added to a round bottom flask containing 100 mL of methanol. Triethylamine (2.4 equiv.), sodium azide (2 equiv.), copper iodide (0.15 equiv.), sodium ascorbate (0.1 equiv.), trans-N-N'-dimethylcyclohexene-1,2-diamine (0.2 equiv.) were sequentially added while purging under nitrogen. The reaction was allowed to stir for 10 mins, and 2-ethynyl pyridine (1 equiv.) was added. Reaction was allowed to stir for 5 mins at room temperature followed by overnight stirring at 55°C. Reaction mixture was filtered over Celite, and the solvent was removed using rotovap. The crude reaction was then purified by liquid chromatography with dichloromethane: ultra (22% MeOH in DCM with 3% NH4OH) 0% to 40% on a 120 gm ISCO silica column.

Z1A34 was synthesized according to the previously reported procedure[1].

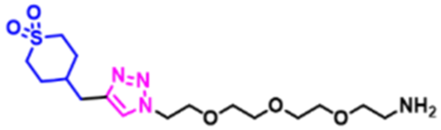

Z1A34

**Z2A19** : Corresponding alkyne: 3-ethynyl thiophene, Peg –linker: Azido-PEG2-amine

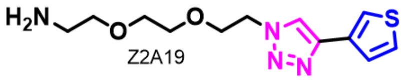

Z2A19

$^{1}$H NMR (600 MHz; $CDCl_3$): 7.8 (s, 1H, aromatic), 7.6 (d, 1H, aromatic), 7.45 (d, 1H, triazole), 7.36 (m, 1H, aromatic), 4.5 (m, 2H, triazoleCH2), 3.89 (m, 2H, triazole CCH2), 3.62 (m, 2H, NCCH2)), 3.58 (m, 2H, methylene), 3.46 (m, 2H, methylene), 2.83 (m, 2H, NH2CH2), 1.45 (s, 2H, amine)

$^{13}$C (600 MHz; $CDCl_3$): 143.8(Cq triazole), 131.9 (Cq- thiophene), 126.2 (CH-triazole), 125.8 (Cq-CH, thiophene), 120.9 (Cq-CH-CH, thiophene), 120.7 (CH, thiophene), 72.7 (NCH2C-), 70.5(-OCCH2O-), 70.1(-OCH2-C-O), 69.5 ( OC-CH2-Ntriazole), 50.3 (-OCH2C-Ntriazole), 41.5 (NH2-C)

ESI: M+1= 283.1

**B2A17**: Corresponding alkyne: 2-ethynyl pyridine, Linker: 3-iodobenzyl amine

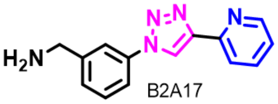

B2A17

$^{1}$H (600 MHz; $CDCl_3$): 8.6 (d, 2H, aromatic), 8.25 (d, 1H, aromatic), 7.84 (s, 1H, triazole), 7.8 (dd, 1H, aromatic), 7.6 (d, 1H, aromatic), 7.5 (t, 1H, aromatic), 7.4 (d, 1H, aromatic), 7.2 (m, 1H, aromatic), 4.0 (s, 2H, NH2-CH2-Ph), 1.54  (broad s, 2H, NH2).

$^{13}$C (600 MHz; CDCl3): 150.01(Cq-aromatic), 149.5 (C-N aromatic), 148.9 (Cq-aromatic), 145.56 (CH aromatic), 137.23 (Cq triazole), 136.99 (CH triazole), 129.94 (CH aromatic),

127.54 (CH aromatic), 123.11 (CH aromatic), 120.45 (CH aromatic),120.03(CH aromatic), 119.15 (CH aromatic), 118.76 (CH aromatic), 46.02 (CH2-NH2)

ESI: M+1 = 252.1545

**Z1A3** synthesis: Corresponding alkyne: 1-bromo-2-ethynylbenzene, Peg linker: Azido-PEG3-amine

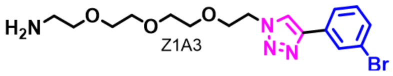

$^1H$ (600 MHz;  $CDCl_3$): 8.3 (s, 1H, triazole), 8.08 (dd, 1H, aromatic), 7.6 (d, 1H, aromatic), 7.39 (t, 1H, aromatic), 7.18 (t, 1H, aromatic), 4.6 (t, 2H, triazoleCH2), 3.9 (t, 2H, triazole CH2-C), 3.6 (s, 4H, methylene), 3.58 (m, 2H, methylene), 3.56 (m, 2H, methylene), 3.4 (m, 2H,methylene), 2.8 (t, 2H, NH2CH2), 2.08 (s, 2H, amine)

$^{13}C$ (600 MHz; $CDCl_3$): 145.22 (Cq triazole) 133.54 (Cq aromatic), 131.42 (CH aromatic), 130.54 (CH aromatic), 129.19 (Cq triazole), 127.63 (CH aromatic), 124.23 (CH aromatic), 121.13 (CH aromatic), 72.33 (N-CH2-C), 70.61 (O-C-CH2), 70.51(O-C-CH2), 70.43 (O-CH2-C), 70.08 (O-CH2-C), 69.49 (Ntriazole-CH2-C), 50.38 (Ntriazole-CH2), 41.3(NH2CH2)

ESI: M+1= 399.1

**Z4A10** : Corresponding alkyne: 1,3 diethynyl benzene, Peg-linker: Azido-PEG4-amine

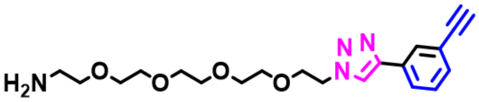

$^1H$ (600 MHz; $CDCl_3$): 8.05 (s, 1H, aromatic), 7.9 (s, 1H, triazole), 7.87 (d, 1H, aromatic), 7.44 (d, 1H, aromatic), 7.38 (t, 1H, aromatic), 4.6 (t, 2H, triazoleCH2), 3.9562 (t, 2H, triazoleCH2-C), 3.6289 (d, 8H, methylene), 3.5814 (d, 4H, methylene), 3.49 (t, 2H, methylene), 3.15 (s, 1H, alkyne), 2.85, (t, 2H, NH2CH2), 2.02 (s, 2H, amine)

$^{13}C$ (600 MHz; $CDCl_3$):   146.8 (C triazole), 131.6 (CH aromatic), 131.2 (CH, aromatic), 129.4 (CH aromatic), 128.9 (C triazole), 126.2 (CH aromatic),122.7 (CH aromatic), 121.4 (C, aromatic), 83.4 (C alkyne), 77.6 (CH alkyne), 73.4(NH2-CH2-C), 70.69(O-CH2-C, methylene), 70.67 (O-CH2-C, methylene), 70.66 (OCH2C, methylene) (O-CH2-C, methylene), 70.63(O-CH2-C), 70.57 (O-C-CH2, methylene), 70.31 (O-C-CH2, methylene), 69.6 (triazole-CH2-C), 50.5 (Ntriazole-$CH_2$), 41.8 ($NH_2$-$CH_2$)

ESI: M+1 = 389.2

**Z1A14**: Corresponding alkyne: 1-ethynyl-2-fluorobenzene, Peg- linker: Azido-PEG3-amine

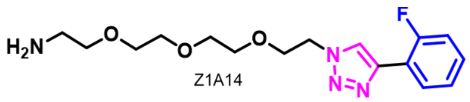

Z1A14

$^{1}$H (600 MHz ; $CDCl_3$) : 8.3 (t, 1H, aromatic), 8.1 (s, 1H, triazole), 7.2 (m, 2H, aromatic), 7.1 (t, 1H, aromatic), 4.6 (s, 2H, triazole-CH2), 3.9 (s, 2H, triazole-CH2-C), 3.6 (m, 8H, methylene), 3.4 (t, 2H, methylene), 2.8 (s, 2H,NH2-CH2), 2.50 (s, 2H, amine)

$^{13}$C (600 MHz; $CDCl_3$): 160.01(Cq aromatic), 159.9 (Cq aromatic split), 158.37 (Cq triazole), 158.34 (Cq triazole split), 141.05 (CH triazole), 141.01(CH triazole split), 129.2 (CH aromatic), 127.7(CH aromatic), 124.5 (CH aromatic), 124.0 (CH aromatic), 118.6 (CH aromatic), 115.5 (CH aromatic), 72.25 (NH2-CH2-C), 72.08 (),70.6 (-OC-CH2), 70.5 (OC-CH2), 70.4 (OCH2-C), 70.1 (O-CH2-C), 69.5 (Ntriazole-CH2-C), 50.3 (Ntriazole-CH2), 41.3 (NH2-CH2)

ESI: M+1 = 339.2

**Z1A16**: Corresponding alkyne: 1-ethynyl-4-fluorobenzene, Peg- linker: Azido-PEG3-amine

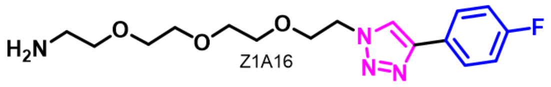

Z1A16

$^{1}$H (600 MHz ; $CDCl_3$) : 7.99 (s, 1H, triazole), 7.8 (m, 2H, aromatic), 7.1(t, 2H, aromatic), 4.6 (t, 2H, triazole CH2), 3.9 (t, 2H, triazoleCH2C), 3.6 (m, 4H, methylene), 3.5 (m, 4H, methylene), 3.48 (m, 2H, methylene), 2.85 (s, 2H, NH2CH2), 2.42 (s, 2H, amine)

$^{13}$C (600 MHz; $CDCl_3$): 163.4 (aromatic), 161.78 (aromatic), 146.79 (C-triazole), 127.46 (aromatic), 127.41(aromatic), 127.03 (aromatic), 127.01 (aromatic), 120.85 (CH-triazole), 115.85 (aromatic), 115.7 (aromatic), 71.97 (NH2-CH2-C), 70.49(-O-C-CH2), 70.47 (O-C-

CH2), 70.39 (O-CH2-C), 70.09 (O-CH2-C), 69.49 (Ntriazole-CH2-C), 50.3 (Ntriazole-CH2), 41.2 (NH2-CH2)

ESI: M+1 = 339.2

**Z1A17**: Corresponding alkyne:  2-ethynyl pyridine, Peg- linker: Azido-PEG3-amine

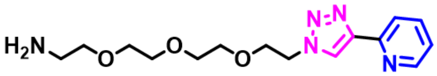

Z1A17

$^{1}$H (600 MHz ; $CDCl_3$) : 8.56 (d, 1H, aromatic), 8.34 (s, 1H, triazole), 8.08 (d, 1H, aromatic), 7.7 (m, 1H, aromatic), 7.2 (m, 1H, aromatic), 4.6 (m, 2H, triazole CH2), 3.93 (m, 2H,triazoleCH2C), 3.63 (s, 4H, methylene), 3.60 (s, 4H, methylene), 3.50 (m, 2H, methylene), 2.89 (m, 2H, CH2NH2), 2.2 ( br s, 2H, amine)

$^{13}$C (600 MHz; $CDCl_3$):  150.4 (Cq, aromatic), 149.3 (CH, aromatic), 148.2 (CH, aromatic), 136.8 (CH, aromatic), 123.2 (CH, aromatic), 122.7 (CH, aromatic), 120.2(CH, aromatic), 72.7 (NH2CH2-C), 70.6 (-OCCH2), 70.5 (-OCCH2), 70.4 (OCH2C), 70.2 (OCH2C), 69.4 (Ntriazole-CH2-C), 50.4 (Ntriazole-CH2), 41.5 (NH2-CH2)

ESI: M+1 = 322.2

**Z4A22**: Corresponding alkyne: 4-(prop-2-yn-1-yloxy)aniline, Peg- linker: Azido-PEG4-amine

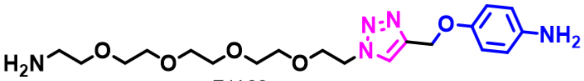

Z4A22

$^{1}$H (600 MHz ; $CDCl_3$) : 7.79 (s, 1H, triazole), 6.8 (d, 2H, aromatic), 6.62 (d, 2H, aromatic), 5.12 (s, 2H, triazoleCH2), 4.56 (m, 2H, triazoleCH2C), 3.87 (m, 2H, methylene), 3.59 (m, 16H, methylene), 2.91 (br s, 2H, NH2CH2), 2.76 ( br s, 2H, amine)

$^{13}$C (600 MHz; $CDCl_3$):  151.4 (aromatic –OCH), 144.4 (aromatic, triazole –CH), 140.57 (aromatic, NH2-CH), 123.9 (aromatic, triazole-C), 116.3113 (aromatic, NH2-C=C-), 116.07 (aromatic, -OC=C-), 72.6 (-OCH2-triazole), 70.58 (NH2-CH2-C-), 70.5 (-OCH2), 70.4 (-

OCH2-C), 70.18 (-OCH2-C), 69.46 (O-CH2), 62.79 (triazole-CH2-C), 50.3(Ntriazole-CH2), 41.5 (NH2-C).

ESI: M+1 = 410.2

**Z4A43**: Corresponding alkyne:  Ethynylcyclopropane, Peg- linker: Azido-PEG4-amine

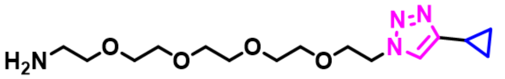

Z4A43

$^{1}H$ (600 MHz ; $CDCl_3$) : 7.4 (s, 1H, triazole), 4.4 (s, 2H, triazoleCH2), 3.76 (m, 2H, triazole CH2C), 3.55 (m, 12H, methylene), 3.47(s, 2H, methylene), 3.16 (s, 2H, methylene), 2.84 (broad s, 2H, amine), 1.85 (m, 1H, cyclic), 0.84 (s, 2H, cyclic), 0.73 (s, 2H, cyclic)

$^{13}C$ (600 MHz; $CDCl_3$): 150.08 (Cq- triazole), 120.9 (CH-triazole), 71.5 (NH2CH2C), 70.5(-OCCH2) 70.47 (-OCCH2), 70.42(OCH2C), 70.09 (OCH2C), 69.6 (triazole-CH2-C), 50.08 (Ntriazole-CH2), 41.1 (NH2-CH2), 7.7 (CH2- cyclic), 6.73 (CH, cyclic).

ESI: M+1 =329.2

**Z1A43**: Corresponding alkyne:  Ethynylcyclopropane, Peg- linker: Azido-PEG3-amine

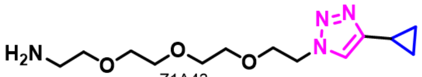

Z1A43

$^{1}H$ (600 MHz ; $CDCl_3$) : 7.4 (s, 1H ,triazole), 4.46 (m, 2H, triazole CH2), 3.8 (m, 2H, triazoleCH2C), 3.58 (s, 8H, methylene), 3.49 (m, 2H, methylene), 2.8 (m, 2H, NH2CH2), 1.9 (m, 1H, cyclic), 1.83 (s, 2H,cmine), 0.9 (m, 2H, cyclic), 0.8 (m, 2H, cyclic)

$^{13}C$ (600 MHz; $CDCl_3$):  149.9(Cq triazole), 120.8(CH triazole), 73.3 (NH2CH2C), 70.5 (-OCCH2), 70.4(-OCH2C), 70.2 (-OCH2C), 69.5 (triazole-CH2-C), 50.0 (Ntriazole-CH2), 41.6 (NH2-CH2), 7.6 (CH2 cyclic), 6.6(CH cyclic)

ESI: M+1 = 285.2

#### NMR and elemental Analysis of Top Three Molecules

Z2A19 –VLVG

$^{1}H$ (600 MHz, D2O): 8.1 (s, 1H, thiophene), 7.7(s, 1H, thiophene), 7.5 (s, 1H, triazole), 7.4 (s, 1H , thiophene),5-3.9 (m, alginate protons),3.6 (s, 2H, methylene), 3.5 (s, 4H, methylene), 3.5- 3.1 (alginate protons), 2.9 (s, 2H, NH2CH2)

Elemental analysis: %C: 36.7, %H: 4.7, %N:7.4

By elemental analysis there is ~26% modification of the alginate backbone.

Z1A3-VLVG:

$^{1}H$ (600 MHz, D2O): 8.4 (s, 1H, triazole), 7.6 (s, 2H, phenyl), 7.3 (s, 1H, phenyl), 7.2 (s, 1H, phenyl), 4.9- 3.8 (m, alginate protons), 3.6 (m, 8H, methylene) 3.4-3.0 (m, alginate protons), 3.06 (2H, t, NH2CH2)

Elemental: %C: 42, %H: 4.9, %N: 4.5

By elemental analysis there is ~19.2% modification of the alginate backbone.

Z4A10-VLVG:

$^{1}H$ (600 MHz, D2O): 8.1 (s, 1H, triazole), 7.5 (s, 2H, phenyl), 7.3 (s, 2H, phenyl), 5.1-3.82 (m, alginate protons), 3.81 (t, 2H, methylene), 3.4 (m, 8H, methylene), 3.4- 3.1 (m, alginate protons), 2.9 (t, 2H, NH2CH2), 2.7 (s, 1H, alkyne)

Elemental: %C: 39.6, %H: 5.37, %N: 4.9

By elemental analysis, there is ~ 25.8% modification of the alginate backbone.

#### Device Manufacturing

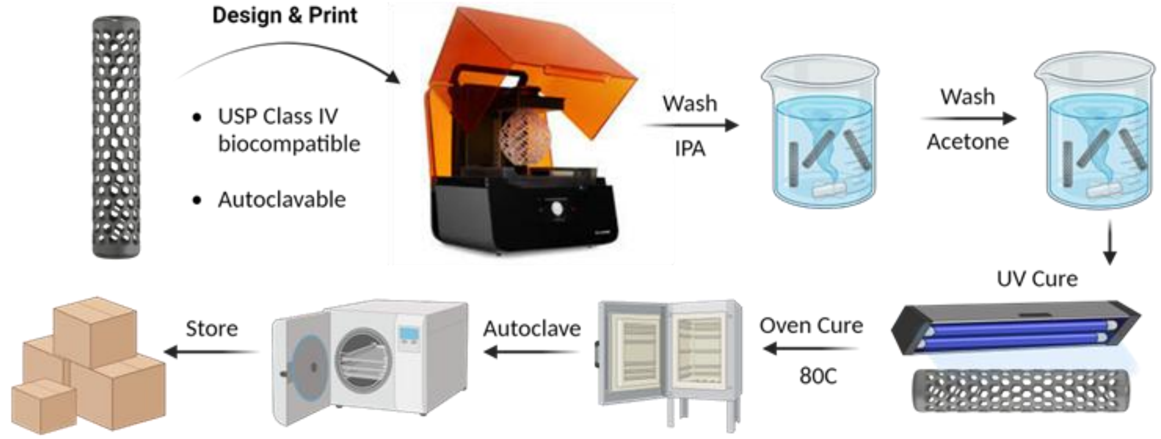

**Supplementary Figure 14. Lattice device fabrication and post-print processing method**. Lattices are 3D printed from Biomedclear resin using a Formlabs Form 4 resin 3D printer. They then undergo several washing and curing steps before final sterilize.
